# Single-Cell Analysis of Human Testis Aging, and Impact of Elevated Body Mass Index

**DOI:** 10.1101/2021.10.19.464550

**Authors:** Xichen Nie, Sarah K. Munyoki, Meena Sukhwani, Nina Schmid, Annika Missel, Benjamin R. Emery, Donor Connect, Jan-Bernd Stukenborg, Artur Mayerhofer, Kyle E. Orwig, Kenneth I. Aston, James M. Hotaling, Bradley R. Cairns, Jingtao Guo

**Author notes:** Correspondence (B.R.C.), (J.G.).

## Abstract

Aging human males display reduced reproductive health, however testis aging is poorly understood at the molecular and genomic level. Here, we utilized single-cell RNA-seq to profile over 44,000 cells from both young and older men (>60 years old) – and examined age-related changes in germline development and in the somatic niche. Interestingly, age-related changes in spermatogonial stem cells appeared modest, whereas age-related dysregulation of spermatogenesis and the somatic niche ranged from moderate to severe. Altered pathways included signaling and inflammation in multiple cell types, metabolic signaling in Sertoli cells, hedgehog signaling and testosterone production in Leydig cells, cell death and growth in testicular peritubular cells, and possible developmental regression in both Leydig and peritubular cells. Remarkably, the extent of dysregulation correlated with body mass index in older, but not younger men. Taken together, we reveal candidate molecular mechanisms underlying the complex testicular changes conferred by aging, and their exacerbation by concurrent chronic conditions such as obesity.

## INTRODUCTION

Parental age has been increasing within industrialized countries (Bray and Gunnell, 2006), and is linked to multiple risk factors including reduced fertility (Harris et al., 2011; Kühnert and Nieschlag, 2004; Levine et al., 2017; NIESCHLAG et al., 1982; Wiener-Megnazi et al., 2012). Aging-associated subfertility is linked to increased time to pregnancy and higher miscarriage rates (du Fossé et al., 2020; Hassan and Killick, 2003). Aging adversely affects sperm genome integrity (DNA fragmentation, telomere length, chromosomal aneuploidy), genome fidelity (DNA mutations), epigenetic status (e.g., DNA methylation), and sperm parameters (e.g., sperm motility, semen volume)(Broer et al., 2013; Cao et al., 2020; Griffin et al., 1996; Kong et al., 2012; Nguyen-Powanda and Robaire, 2020; Pohl et al., 2021; Sharma et al., 2015; Singh et al., 2003). Aging also impacts the entire endocrine system, including the hypothalamic–pituitary–gonadal (HPG) axis which is critical for the production of hormones such as testosterone (T) and leads to changes in testis physiology as well as fertility status (Decaroli and Rochira, 2016; Pincus et al., 1996; Sampson et al., 2007; Wu et al., 2008). Although histological studies have revealed aging-associated testicular changes in both humans and mice (Gosden et al., 1982; Santiago et al., 2019), it is still largely unknown at the molecular and genomic level how aging separately impacts the human germline and the supporting cells of the testis. Progress has been slowed by the difficulty in obtaining sufficient testis samples (both young/healthy and older) for in-depth studies, and the difficulty in distinguishing between the effects of aging in relation to possible concurrent factors, such as increased body mass index (BMI) and tobacco smoking status. Thus, a detailed molecular understanding of aging-related changes requires well annotated samples in sufficient numbers, and the application of new molecular and genomics tools to provide systematic and mechanistic insights.

Human spermatogenesis involves the differentiation of spermatogonial stem cells (SSC) into mature sperm through a complex developmental process (de Kretser et al., 1998). As the only germline stem cell, SSCs must carefully balance their self-renewal and differentiation, and then undergo niche-guided transitions between multiple cell states and cellular processes – including an initial commitment to mitotic growth/proliferation, followed by meiosis and the subsequent stages of sperm maturation (Neto et al., 2016). The testicular microenvironment is composed of multiple somatic cell types, including Sertoli cells, Leydig cells and testicular peritubular cells (TPCs; also called peritubular myoid cells). These cells provide physical and biochemical interactions and signaling among themselves and with the germ cells along multiple stages of germ cell development – to ensure that spermatogenesis is continuously executed (Oatley and Brinster, 2012).

The integrity of testis physiology is critical for successful sperm production (Oatley and Brinster, 2012). Previous histomorphological studies of aging men have demonstrated a decrease in Leydig and Sertoli cells, and in some samples a thickening of the basement membrane(Dakouane et al., 2005; Jiang et al., 2014; Neaves et al., 1984; Santiago et al., 2019), but the molecular alternations that underlie these changes remain unknown. However, many healthy fertile older men maintain a relatively normal testis physiology and comparable number of round spermatids (Pohl et al., 2019). Thus, both the basis for the heterogeneity observed in populations, and the molecular mechanisms underlying the physiological and histological changes during testis aging remain elusive.

Recent advances in single cell genomics enable the examination of thousands of individual cells, including those of entire organs without the need for prior sorting or enrichment procedures. Work from several groups has proven the power of single cell RNA-seq (scRNA-seq) profiling to study human testis development and pathology (Guo et al., 2017, 2018, 2020, 2021; Sohni et al., 2019; Shami et al., 2020; Wang and Xu, 2020). Here, we continue to leverage our unique sample resource (involving rapid autopsy testis samples from cadaveric organ donors) to apply scRNA-seq and other approaches such as immunostaining. We profiled single cell transcriptomes of testes from all eight older adults (>60 years old) in our collection, and compared to four young fertile adults (17-22 years old) – resulting in a dataset consisting of over 44,000 single cell transcriptomes. We examined the transcriptional changes within each major testicular cell type during aging, which revealed a set of molecular mechanisms underlying human testis aging. Remarkably, the data partitioned older donors according to their BMI status, and pronounced dysregulation was observed in older donors with high BMI (>30). To explore further, we validated the genomics results by conducting extensive immunohistostaining as well as initial functional experiments. Taken together, our study 1) provides major insights into how aging might combine with other concurrent factors (such as BMI) to impact germ cells and their niche in the human testis, 2) reveals potential biomarkers for diagnosis of testis aging and directions for potential treatment of aging-related subfertility, and 3) serves as a foundational dataset for the scientific community to study how human testis and fertility respond to aging and elevated BMI.

## RESULTS

### Single-cell transcriptomes of testes of young and older men

To investigate human testis aging, we utilized from our rapid autopsy collection 8 whole testes from donors aged 62 to 76 years old (termed ‘older’) and 4 whole testes from healthy donors aged 17 to 22 years old (termed ‘young’; **Fig1A**). As the focus of our current study involves the normal physiological process of aging rather than diseased states (e.g., infertility), we confirmed that all older donors we examined had offspring, indicting they lacked major fertility issues while they were young. Histologically, we observed (via PAS staining) complete spermatogenesis in all of the younger testes, and also in the majority (5) of the older testes. However, the testes from three older males had apparent impaired spermatogenesis (**Fig 1B**). In addition, we found an elevated thickness of walls of seminiferous tubules (**Fig 1B**) and an increase in extracellular matrix (ECM) deposition in interstitial tissues of all older testes compared to young (**Fig S1B**), consistent with a previous study (Johnson et al., 1986).

**Figure 1.**
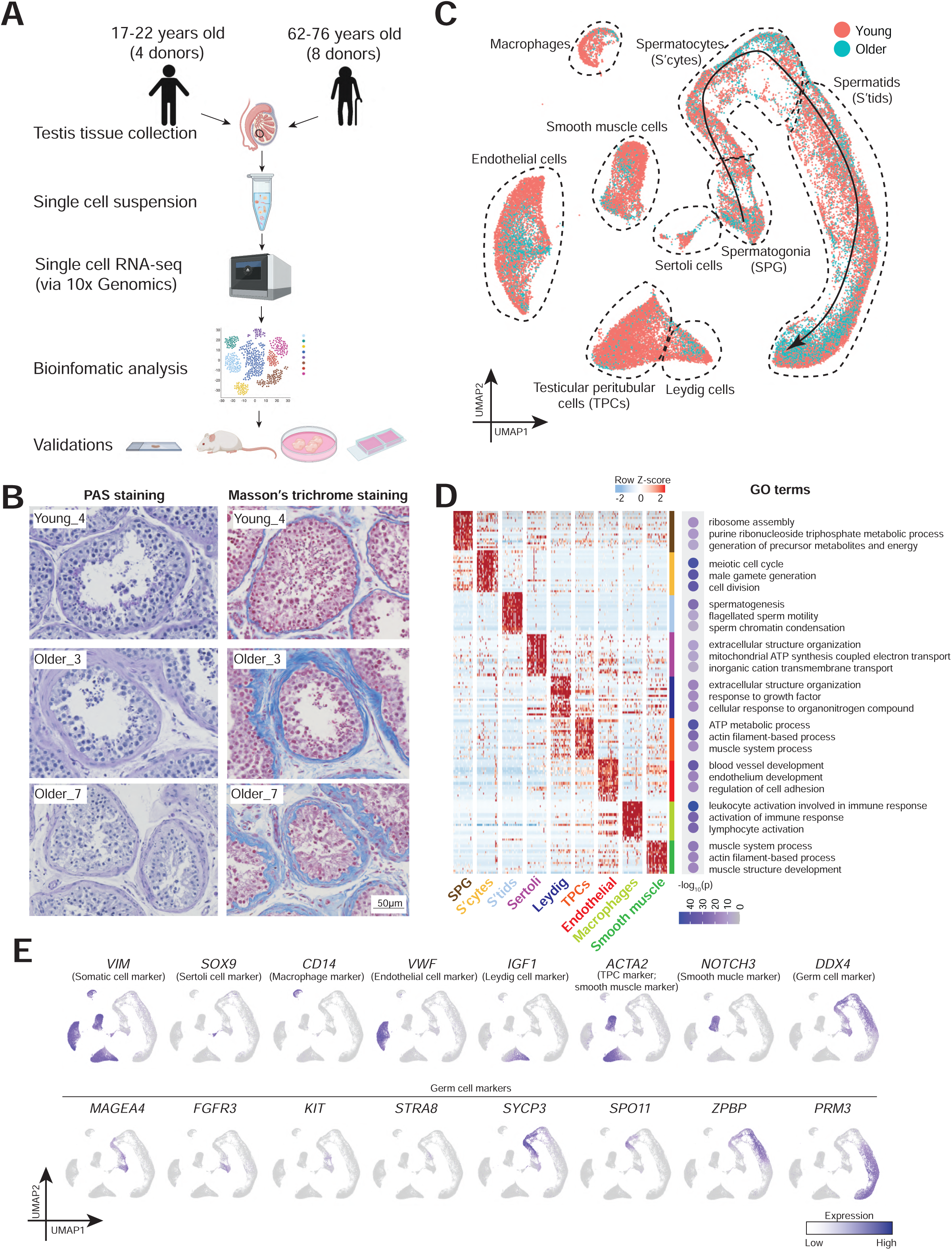
Single cell transcriptome profile of testicular cells from young and older men. (A) Schematic of the experimental workflow. (B) Periodic acid–Schiff (PAS) staining (left) and Masson’s trichrome staining (right) of sections of testis from young and older donors, respectively, demonstrating morphological changes during aging and variations among older human testis. (C) UMAP plot showing the annotated testicular cell types from both young and older men (n=12). Each dot represents a single testicular cell, and is colored based on its donor of origin. (D) Left: Heatmap showing the top 200 differentially expressed genes of each cell cluster from (C). The scaled gene expression levels are colored according to Z-score. Right: The corresponding top three GO terms enriched in the marker genes of each cell cluster with -log10(P-value) colored according to the color key at the bottom. (E) Expression of selected markers identifying major testicular cell types cast on the UMAP plot. Purple (or grey) represents a high (or low) expression level as shown on the color key at the right bottom.

To understand the molecular alterations that accompanied aging, we performed scRNA-seq on all 12 samples using the 10x Genomics Chromium platform. A total of 44,657 cells (13,156 cells from young, 31,501 cells from older) passed quality control and were analyzed, yielding ∼ 2600 genes/cell (**Fig S1A**). To identify cell clusters via Uniform Manifold Approximation and Projection (UMAP), we projected expression of sets of known cell type-specific marker genes and identified 6 major somatic cell types, along with germ cells at various phases, including: Sertoli cells (*SOX9*^+^), macrophages (*CD14*^+^), endothelial cells (*VWF*^+^), Leydig cells (*IGF1*^+^), TPCs (*ACTA2*^+^ and *WFDC1*^+^), smooth muscle cells (*NOTCH3*^+^), undifferentiated and differentiating spermatogonia (*FGFR3*^+^ or *KIT*^+^), spermatocytes (*SYCP3*^+^), and post-meiotic spermatids (*PRM3*^+^) (**Fig 1C** and **1E**; see **Fig S1D** for additional markers/examples). The assigned cell types were further confirmed by Gene Ontology (GO) analysis of differentially expressed genes (DEGs) (**Fig 1D**). Notably, initial comparisons suggested heterogeneity in the older testis samples (**Fig 1C** and **Fig S1C**), requiring detailed analyses to understand their heterogeneity and identify age-related differences, conducted below.

### Analysis of aging in spermatogonia

To understand how the germline changes during aging, we performed a focused analysis of germ cell clusters from Fig 1C. We classified 7 phases of germ cell development based on the following marker genes: spermatogonial stem cells (SSCs; *PIWIL4*^+^ or *GFRA1*^+^), differentiating spermatogonia (*KIT*^+^ and largely *MKI67*^+^), early primary spermatocytes (from *STRA8*+ to *SPO11*+), late primary spermatocytes (*SPAG6*^+^/*ZPBP*^+^/*ACRV1*^-^), round spermatids (*ACRV1+*), elongating spermatids and elongated spermatids (*PRM3*^+^) (**Fig 2A-B**). We further examined the distribution of cell types in each donor sample and quantified their relative proportions at different stages of spermatogenesis (**Fig 2C-D**). Complete spermatogenesis (including typical cell type proportions) was observed in 5 of 8 older donors, whereas varying degrees of spermatogenesis impairment were observed in the remaining 3 of 8 older donors, in agreement with our histological examination. However, with all 3 of the testis samples from older donors undifferentiated SSCs were still observed in moderate numbers, whereas germ cell types further in development were highly diminished, indicating that germ cell differentiation was heterogeneously impacted by aging, consistent with previous histological examinations (Paniagua et al., 1991).

**Figure 2.**
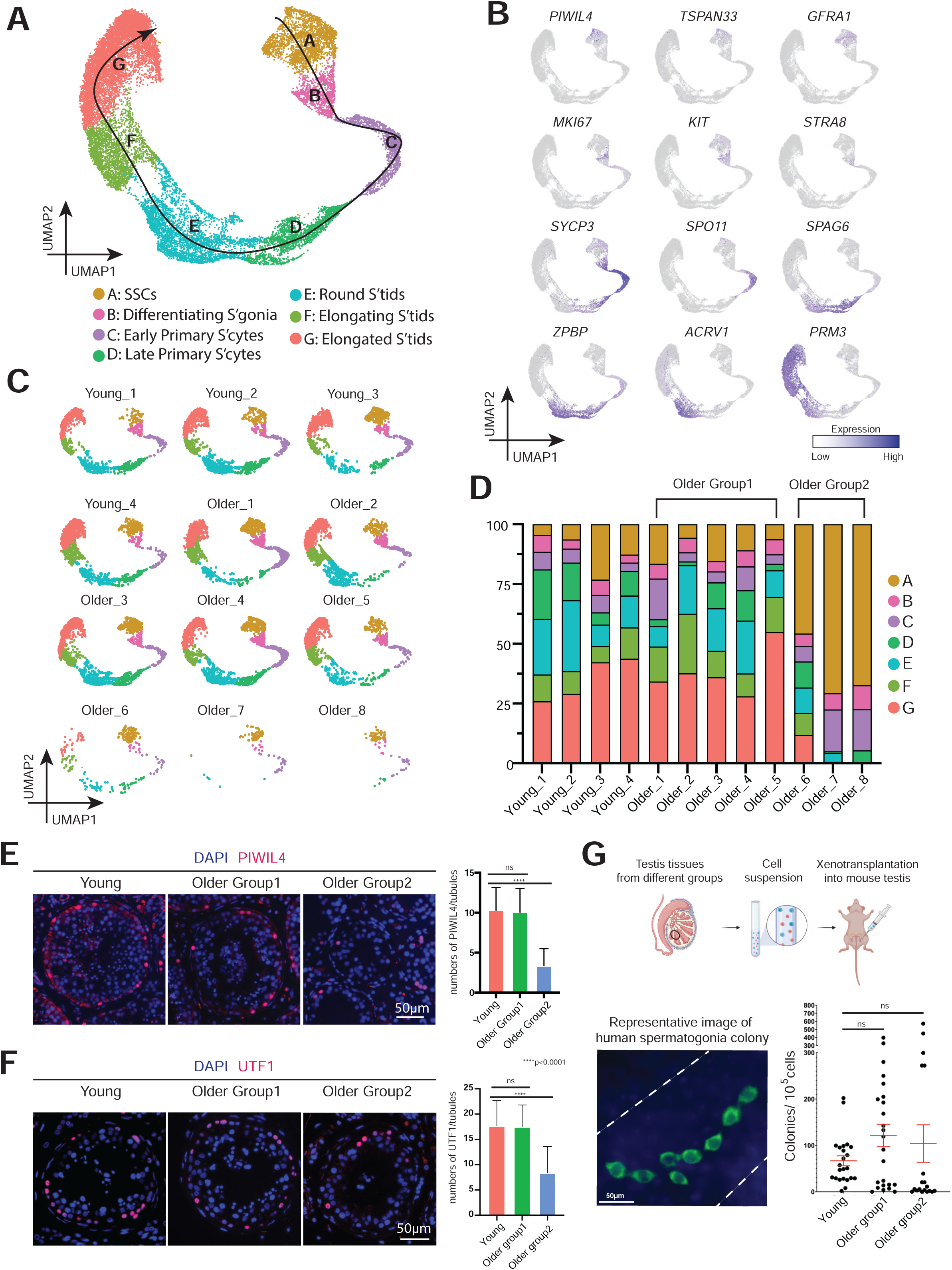
Strong variation in spermatogenic dynamics observed in men with advanced age. (A) UMAP plot showing the annotated germ cells captured from Fig 1C. (B) Expression patterns of selected markers cast on the UMAP plot in Fig 2A. Purple (or grey) represents a high (or low) expression level as shown on the color key at the right bottom. (C) Deconvolution of the UMAP plot of Fig 2A according to donors of origin. (D) Bar plot showing the percentage of major germ cell types for each individual. Older donors with complete spermatogenesis are classified as Older Group1, while older donors with spermatogenic impairment are classified as Older Group2. (E) Left: Immunofluorescence image of a State 0 marker, PIWIL4 (red) across three donor groups. Nuclei were counterstained with DAPI (blue). Right: Quantification of the number of PIWIL4-positive cells in cross-section of each seminiferous tubule in different groups. Bars represent mean with standard deviation (SD) of 20 independent tubules per group. n = 6 human samples. ns, no significance. ****p < 0.0001 (two-tailed t-test). (F) Left: Immunofluorescence image of a State 1 marker, UTF1 (red) across three donor groups. Nuclei were counterstained with DAPI (blue). Right: Quantification of the number of UTF1-positive cells in cross-section of each seminiferous tubule in different groups. Bars represent mean with SD of 20 independent tubules per group. n = 6 human samples. ns, no significance. ****p < 0.0001 (two-tailed t-test). (G) Top: Schematic of the human to nude mouse xenotransplantation experiment. Bottom left: Immunofluorescence images of whole-mount staining of recipient mouse testes, using an anti-primate antibody. Human spermatogonia colonies were observed after two months of xenotransplant. Scale bar, 50 µm. Bottom right: Quantification of the number of colonies per 10^5^ viable cells transplanted per testis in different groups. Bars represent mean with SD. n = 9 human samples.

To further explore potential molecular mechanisms underlying testis aging and aging-induced male subfertility, we separated older donors into two groups: donors with histologically normal spermatogenesis were classified as Older Group1, and donors with clearly defective spermatogenesis were classified as Older Group2. Here, we hypothesized that if certain aging-associated changes lead to male subfertility, we should observe (compared to young samples) a modestly dysregulated transcriptional signature within Older Group1, which becomes stronger in Older Group2.

We then reanalyzed spermatogonia clusters from Fig 2A (**Fig S2A-B**). As observed previously, spermatogonia partitioned into 5 cellular states (**Fig S2A**). However, we did not observe DEGs that exceeded our statistical cutoffs across the three groups (Young, Older Group1, and Older Group2) (**Fig S2C-D**). Next, we identified DEGs across these different groups during spermatogenesis, which only emerged at/after the elongated spermatid stage when comparing Young and Older Group1 (Note: Older Group2 lacked sufficient elongated spermatids to analyze). Here, GO analysis in older elongated spermatids revealed upregulated DEGs enriched in categories for protein targeting, while downregulated DEGs enriched categories for peptide chain elongation and oxidative phosphorylation (**Fig S2E-F**).

Regarding SSC abundance, both the numbers of State 0 and State1 cells (calculated by PIWIL4 or UTF1 staining) did not change between Young and Older Group1, but declined significantly in Older Group2, indicating that aging-associated spermatogenesis deficiency is due at least in part to the quantity of SSCs (**Fig 2E-F** and **S2G-H**). To further assess the potential activity of older SSCs compared to that of young SSCs, we conducted SSC xenotransplantation experiments (Brinster and Zimmermann, 1994; Valli et al., 2014). Here, heterogenous cell suspension generated from human donor testis tissue was injected into the testis of a nude mouse that was rendered infertile by prior busulfan treatment. Notably, two months after xenotransplantation, we found an overall comparable average transplant efficiency across the three Groups (**Fig 2G**), which is consistent with the scRNA-seq result that age-associated transcriptomic change of SSCs is subtle. Interestingly, we noticed higher variation in colony forming potential in the Older Groups compared to the younger group. Taken together, these results indicate that aging spermatogonia show limited transcriptional changes and retain developmental potential, while displaying higher variability.

### Global alterations of niche cells and niche-germline interactions in the aging human testis

To understand how the SSC niche changes during aging, we identified DEGs in each somatic cell type (**Fig 3A**). Changes of individual genes within specific cell types are detailed later, whereas we will explore here the global trends and GO categories that accompany aging across multiple cell types. Most DEGs were cell-type specific (upregulated: 367; downregulated: 683), however, a subset of upregulated genes were shared among two or more somatic cell types (25 genes shared in more than 3 cell types and 139 genes shared in 2 or 3 cell types), suggesting commonality. We detected inflammation-related GO terms including “cytokine signaling in immune system” as a hallmark of aging (**Fig 3A**). Example genes are listed, including *TIMP1* (shared by 5 somatic cell types), a secreted protein in inflammatory networks (Knight et al., 2019), *IL6ST* (shared by 3 somatic cell types), the *IL6* receptor, and *IFITM2/3* (shared by 3 somatic cell types), representing genes induced by interferon (**Fig 3B**). Notably, their expression progressively changed from Young to Older Group1 to Older Group2, supporting our hypothesis that the aging-associated genes which are involved in subfertility display moderate dysregulation in Older Group1, and stronger dysregulation in Older Group2. For validation, we stained TIMP1 (**Fig 3C** and **S3F**), which is secreted by Sertoli cells inside seminiferous tubules, and we found its signal elevated from Young to Older Group1 to Older Group2. GO analysis for shared downregulated genes was limited to changes shared between Leydig cells and TPCs rather than the whole microenvironment, since there were no shared downregulated genes between more than 2 cell types. Collectively, the results suggest inflammation as a common feature of aging testis somatic cells.

**Figure 3.**
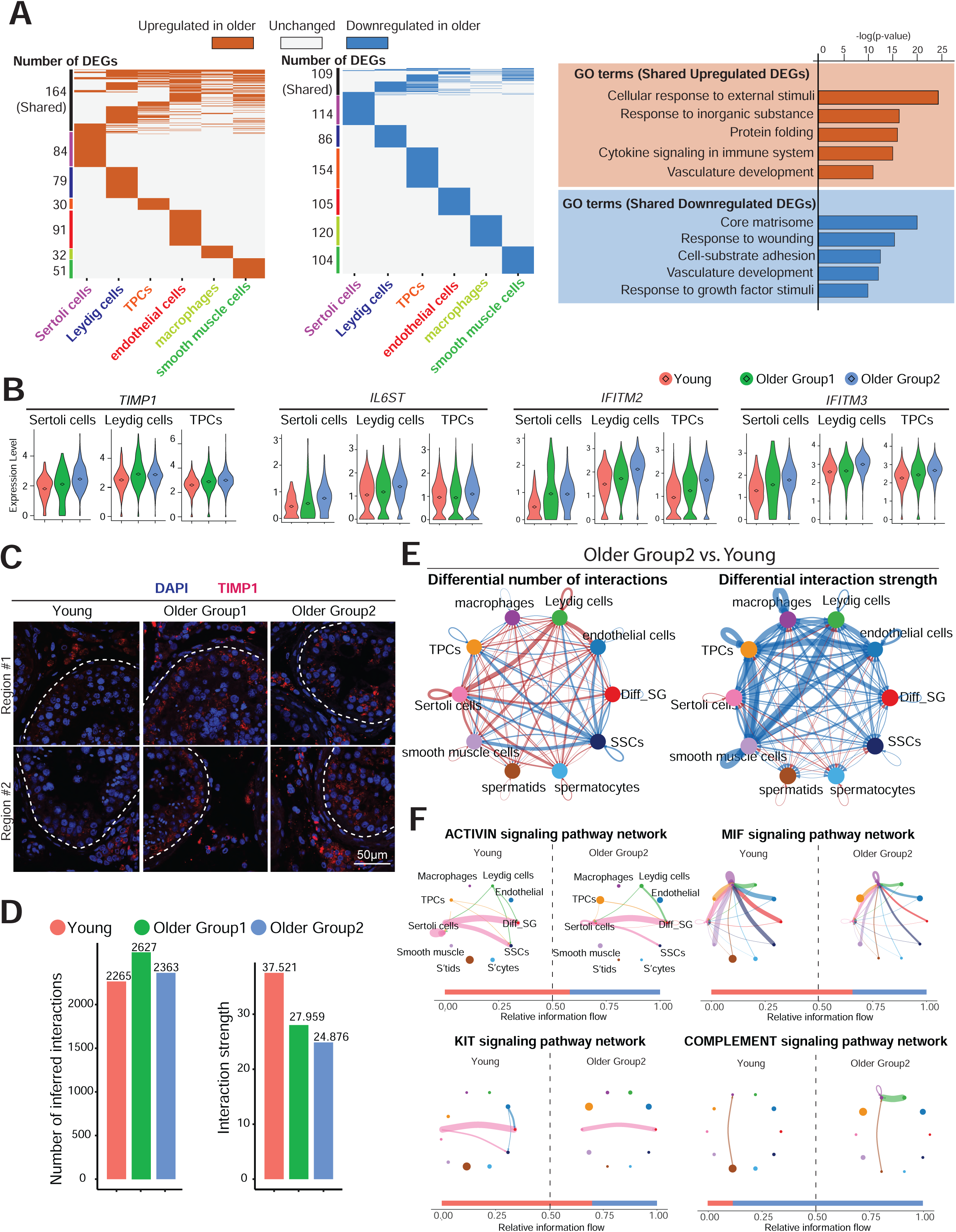
Global alterations of niche cells and niche-germline interactions in older human testis. (A) Left: Heatmaps showing common and unique upregulated DEGs between young and older men in each major somatic niche cell type. Middle: Heatmaps showing common and unique downregulated DEGs between young and older men in each major niche cell type. Right: Representative GO terms of shared upregulated (top) or downregulated (bottom) DEGs between young and older testis niche cells and their associated p-value. (B) Violin plots showing upregulated DEGs of older niche cells related to inflammation. Diamond inside the violin represents mean. (C) Immunofluorescence images of TIMP1 (red) show the elevated TIMP1 staining signal in older testis. Nuclei were counterstained with DAPI (blue). Dashed lines indicate seminiferous tubule membranes. Two donors from each group are shown. Scale bar, 50 µm. (D) Bar plots showing the number of inferred interactions (left) or interaction strength (right) in the cell-cell communication network analyzed by CellChat across three groups. (E) Circle plots (by CellChat analysis) depict the differential number of interactions (left) or interaction strength (right) in the cell-cell communication network between Young and Older Group2, respectively, indicating SSCs lose interactions with niche cells during aging. Red or blue edges represent increased or decreased signaling in Older Group2 compared to Young, respectively. (F) Circle plots showing selected inferred differential signaling networks. The edge width represents the communication probability. Bar graph at the bottom of each panel illustrating representative information flow in Older Group2 (blue) and Young (red).

We then performed CellChat analysis to investigate the niche-germline interactions. Interaction numbers did not change significantly between younger and older testes, but interaction strength weakened from Young to Older Group1 to Older Group2 (**Fig 3D**). In particular, SSCs displayed lower interaction numbers and interaction strength with somatic cells in Older Group2 compared with Young (**Fig 3E**). For example, the activin signaling pathway between Sertoli cells (senders) and SSCs (receivers) decreased in Older Group2. Notably, Activin signaling may help regulate SSC self-renewal and differentiation (**Fig 3F** and **S3B**), consistent with prior work(Guo et al., 2020).Furthermore, the KIT signaling pathway decreased between Sertoli cells (senders) and differentiating spermatogonia (receivers), a pathway also known to play a key role in maintaining the balance of SSCs self-renewal and differentiation (**Fig 3F** and **S3C**) (Schrans-Stassen et al., 1999; Rossi et al., 2000). In addition, signaling pathways related to the immune system also changed during aging, including a decrease in the MIF signaling pathway and an increase in the complement signaling pathway in macrophages in Older Group2 (**Fig 3F** and **S3D-E**). The trend of these pathways was also true in Older Group1 compared to Young (**Fig S3A**). Taken together, these results suggest that niche cell signaling changes during aging in a manner detrimental to germline development.

### Sertoli cells display inflammation and metabolic dysregulation during aging

To explore Sertoli cell alterations during aging, we performed a focused analysis of the Sertoli cell cluster from Fig 1C and arranged the cells along pseudotime development trajectory (**Fig 4A-B**). The deconvoluted pseudotime plot showed Sertoli cells from Young samples residing at the beginning of the trajectory, cells from Older Group2 settling at the end of the trajectory, and cells from Older Group1 distributed along the trajectory (**Fig 4B-C**), indicating that age-dependent changes in Sertoli cells exhibited by Older Group2 may represent more pronounced changes along the same pathway(s) as Older Group1.

**Figure 4.**
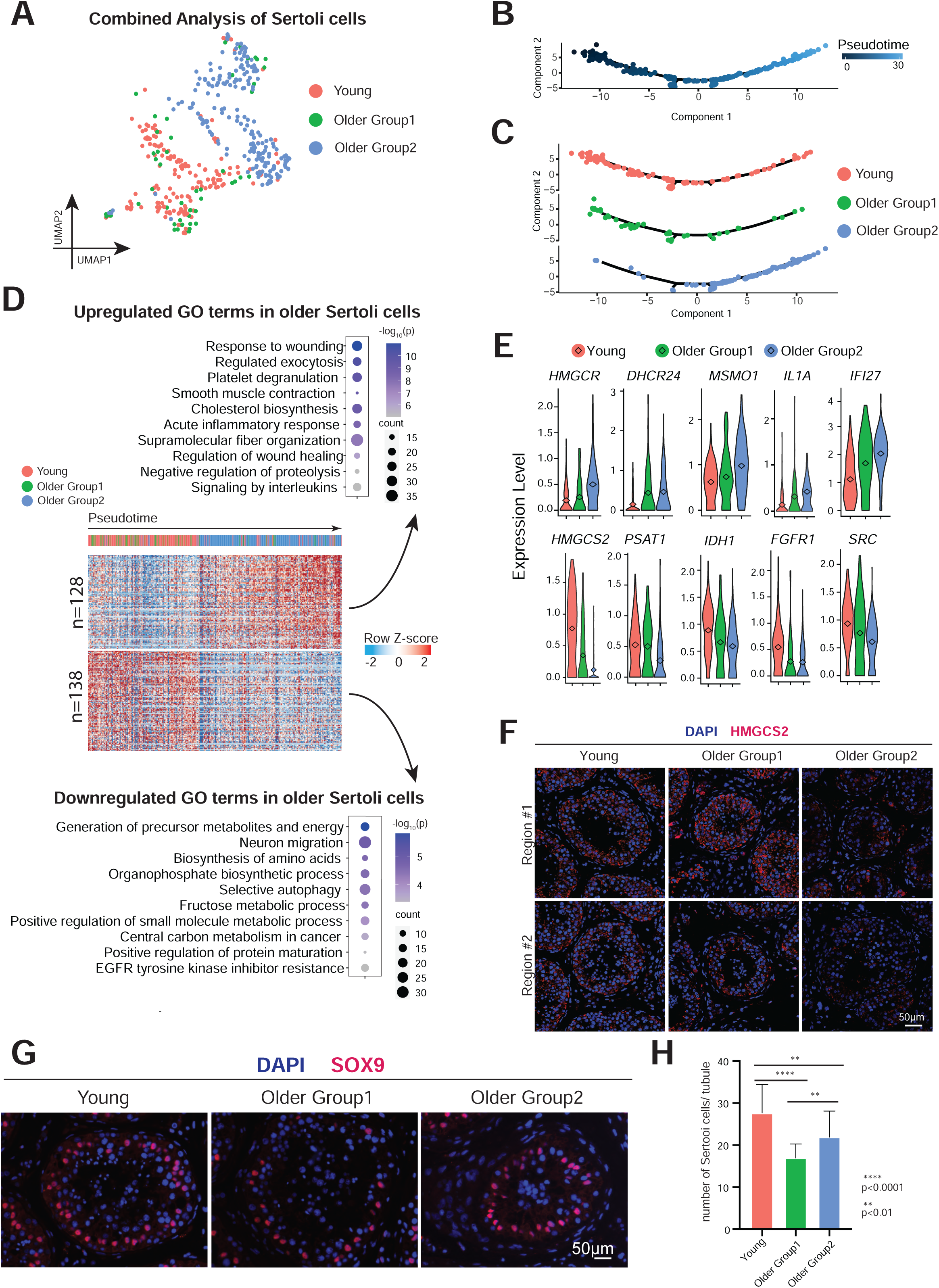
Sertoli cells from older men display metabolic dysregulation. (A) UMAP plot showing focused analysis of Sertoli cells from Fig 1C. (B) Pseudotime trajectory of Sertoli cells analyzed by Monocle. (C) Deconvolution of pseudotime trajectory of Fig 4B according to Young/Older Groups, predicting gradual changing from Young through Older Group1 to Older Group2. (D) Heatmap showing both upregulated and downregulated DEGs in Older Groups of Sertoli cells with columns/cells placed in Pseudotime order defined in Figure 4C. The scaled gene expression levels are colored according to Z-score. The top 10 upregulated or downregulated GO terms enriched in the DEGs are listed with P-value and gene numbers. (E) Violin plots showing upregulated DEGs of older Sertoli cells on the top panel and downregulated DEGs of older Sertoli cells at the bottom panel. The diamond inside the violin plot represents the mean. (F) Immunofluorescence images of HMGCS2 (red) in different groups show significant decrease in staining signal in Older Group2 testis. Nuclei were counterstained with DAPI (blue). Two donors from each group are shown. (G) Immunofluorescence images of a Sertoli cell marker, SOX9 (red) in different groups (Young, Older Group1, and Older Group2). Nuclei were counterstained with DAPI (blue). (H) Quantification of the number of Sertoli cells (SOX9 positive) in cross-section of each seminiferous tubule in different groups. Bars represent mean with SD of 20 independent tubules per group. n = 6 human samples. ****p < 0.0001, **p < 0.01 (two-tailed t-test).

Combined analysis of pseudotime with differential gene expression revealed 128 upregulated genes and 138 downregulated genes during Sertoli cell aging (**Fig 4D**). The upregulated genes in older Sertoli cells are enriched for inflammation-related GO terms, like “response to wounding”, “acute inflammatory response”, and “signaling by interleukins” (including *IL1A, IFI27, MIF, IL6ST*, and *CCL2*), consistent with our global analysis of niche cells (**Fig 4D-E** and **S4A-B**). Interestingly, upregulated GO terms included cholesterol biosynthesis, which included enzymes functioning in lipid metabolism upstream of cholesterol biosynthesis (e.g., *HMGCR*), enzymes directly involved in cholesterol biosynthesis (e.g., *DHCR24, MSMO1*, and *SQLE*), and translocator proteins or enzymes that are downstream of cholesterol pathway (e.g., *TSPO, FADS1*, and *ELOVL5*) (**Fig 4D-E** and **S4A-B**). Our observations may explain the mechanism underlying prior work showing that older human Sertoli cells accumulate lipid droplets in their cytoplasm (Paniagua et al., 1991). The downregulated genes were enriched for GO terms related to metabolism, like “generation of precursor metabolites and energy” (e.g., *HMGCS2, IDH1, FGFR1, SRC, ACO2, UQCRC1, AKR1B1*, and *PFKFB4*), “biosynthesis of amino acids” (e.g., *PSAT1*), “organophosphate biosynthetic process”, “positive regulation of small molecule metabolism”, and “positive regulation of protein maturation” (**Fig 4D-E** and **S4A-B**). To confirm the gene expression pattern, we performed an immunofluorescence (IF) staining as well as a Western blot of HMGCS2 in all three groups and found Older Group2 showed a significant decrease of HMGCS2 in Sertoli cells (**Fig 4F** and **S4C-D**). This trend also held true for PSAT1 in Western blot analysis, which was only expressed in Sertoli cells, reinforcing our scRNA-seq data (**Fig S4D**). Furthermore, Western blot analysis of IDH1 confirmed its gradual decline in older groups (**Fig S4D**). As germ cells receive nutrients from Sertoli cells for spermatogenesis, our results suggest that alterations in metabolic behavior in aging Sertoli cells likely contribute to spermatogenic failure (Petersen and Söder, 2006).

We next assessed Sertoli cell numbers by staining for SOX9, a known Sertoli cell marker localized to the nucleus (**Fig 4G-H** and **S4E**). Sertoli cell numbers declined significantly in both Older Group1 and Older Group2 compared to Young, aligned with previous reports (Santiago et al., 2019). However, the decline in Older Group2 was not as pronounced as that of Older Group1, likely due to vertical/diameter regression of seminiferous tubules in Older Group2. Collectively, the data suggest that both loss of Sertoli cells and diminishing metabolic function may influence spermatogenesis in males with advanced age.

### Leydig cells display dysregulation in signaling, testosterone and developmental identity

A focused analysis of the Leydig cell cluster from Fig 1C yielded 174 genes sequentially upregulated and 144 genes sequentially downregulated from Young to Older Group1 to Older Group2 (**Fig 5A-B**). The GO terms enriched in the upregulated genes of older Leydig cells were largely associated with smooth muscle contraction (e.g., *ACTA2, MYH11, TPM1/2, MYL9*, and *FLNA*) (**Fig 5B-C** and **S5A**), growth suppression, and reactive oxygen species (ROS). Notably, smooth muscle contraction is a feature of TPCs, which develop from the same cell progenitor as Leydig cells during puberty (Guo et al., 2020), indicating that aging Leydig cells partially lose their cell identity and acquire transcriptome features of TPCs. Regarding growth, older Leydig cells upregulate *PTEN, RHOB*, and *ROCK1/2*, which suppress cell survival and proliferation (**Fig S5A** Regarding ROS, older Leydig cells upregulate multiple genes, including *PRDX6, SOD2, MT2A, MT1X, NAMPT*, and *HIF1A* (**Fig 5B-C** and **S5A**). These findings were further confirmed by Gene Set Enrichment Analysis (GSEA) which focuses on gene sets that share common biological functions (**Fig S5C**).

**Figure 5.**
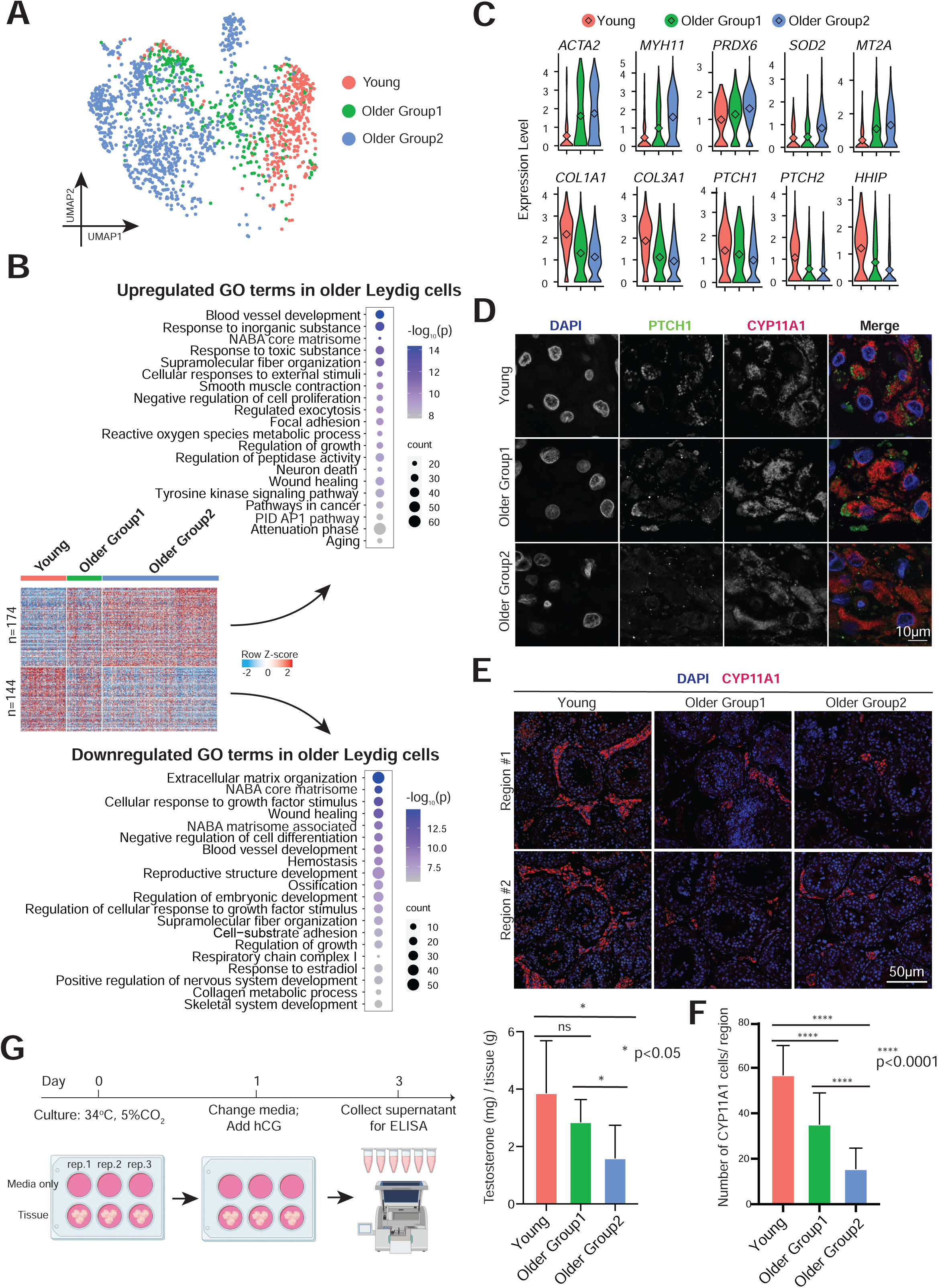
Leydig cells from older men display declined functionality, including lower production of testosterone. (A) UMAP plot showing focused analysis of Leydig cells from Fig 1C. (B) Heatmap showing both upregulated and downregulated DEGs in older Leydig cells. The scaled gene expression levels are colored according to Z-score. The top 20 upregulated or downregulated GO terms enriched in the DEGs are listed with P-value and gene numbers. (C) Violin plots showing upregulated DEGs of older Leydig cells within the top panel and downregulated DEGs of older Leydig cells within the bottom panel. The diamond inside the violin plot represents mean. (D) Immunofluorescence images of PTCH1 (green) with a Leydig cell marker, CYP11A1 (red) in different groups, revealing decreased PTCH1 expression in older Leydig cells. Nuclei were counterstained with DAPI (blue). Two donors from each group are shown. (E) Immunofluorescence images of a Leydig cell marker, CYP11A1 (red) in different groups. Nuclei were counterstained with DAPI (blue). Two donors from each group are shown. (F) Quantification of the number of Leydig cells (CYP11A1 positive) in cross-section of each microscopic field (0.04 mm^2^) in different groups, demonstrating reduced Leydig cells during aging. Bars represent mean with SD of 20 independent tubules per group. n = 6 human samples. ****p < 0.0001(two-tailed t test). (G) Left: Schematic of tissue explant culture used to test testosterone production of older Leydig cells. Right: Quantification of testosterone production in Young, Older Group1, and Older Group2, indicating older testis produce less testosterone. Bars represent mean with SD of 9 donors with 3 technical replicates per sample. *p < 0.05, ns: no significance (two-tailed t-test).

Leydig cells create ECM components, but downregulated genes for ECM organization during aging (e.g., *COL1A1/2, COL3A1, COL5A1/2*), as well as many genes encoding secreted proteins with diverse functions such as *VIT, SFRP1, IGF2, PENK*, and *LTBP4* (**Fig 5B-C** and **S5B**). GO analysis indicated that older Leydig cells may respond less well to growth factor stimuli, as downregulated membrane receptors included *PDGFRA, ITM2A, SCARA5, SPRY1, ROR1, FGFR1*, and *EGFR* (**Fig S5B**). In particular, key components of Hedgehog (HH) signaling pathway were downregulated in older Leydig cells, including *PTCH1/2* and *HHIP* (**Fig 5C**). CellChat analysis further suggested HH signaling pathway dysregulation in Leydig cells (**Fig S5D**). To validate, we co-stained for PTCH1 and the Leydig cell marker CYP11A1. Consistently, the protein level of PTCH1 decreased from Young to Older Group1 to Older Group2 (**Fig 5D** and **S5E**). Taken together, these results suggest that aging Leydig cells likely undergo both intrinsic and extrinsic dysregulation of pathways involved in key signaling pathways and cell survival.

Next, we quantified Leydig cell numbers across the three groups (Young, Older Group1, Older Group2) through IF staining of CYP11A1, and observed a significant decrease during aging (**Fig 5E-F**). For functional validation, we assessed the ability of older Leydig cells to produce testosterone. Here, we cultured the same weight of small pieces of testicular tissue *in vitro*. We replaced culture media at 24 hours with media supplemented with human chorionic gonadotropin (hCG), a hormone that stimulates testosterone production, and we evaluated testosterone concentration in the supernatant 48h after addition of the hCG-supplemented media (**Fig 5G**). Notably, testosterone declined significantly in Older Group2 compared to Young. Here, we note that the same weight of tissues contains fewer Leydig cells during aging, as our staining indicated. Altogether, we conclude that the remaining Leydig cells in older testes produce less testosterone per unit volume of seminiferous tubule in response to hCG stimulus.

### Aging testicular peritubular cells show dysregulated cell death, matrisome, developmental pathways and contractility

We next analyzed the TPC cluster from Fig 1C, examined age-associated changes shared between the two older groups of TPCs compared to young TPCs (**Fig 6A**), and as previewed above, observed dysregulation of inflammation in older samples (**Fig 6B**). Notably, older TPCs were enriched in “positive regulation of cell death”, and displayed increased transcription of *CDKN1A, MDM2, ZMAT3, MEG3*, and *GADD45A*, all of which are involved in p53-directed cell cycle arrest or cell death (**Fig 6B-C** and **S6B**). IF staining of p21 (encoded by *CDKN1A*) and ACTA2 (an TPC marker) revealed clear increases in the number of p21 positive TPC cells in Older Group1 and Older Group2 compared to the Young group, consistent with our scRNA-seq data (**Fig 6D**). Interestingly, the number of TPCs that surround older seminiferous tubules significantly increased as well, indicating the presence of unknown (possibly post-transcriptional) mechanisms that underlie the survival and proliferation of older TPCs (**Fig 6I**). The diameters of seminiferous tubules were largely the same in Young and Older Group1, but decreased in Older Groups2 (**Fig 6J**). Intriguingly, GO analysis indicates the upregulation of autophagy in older TPCs (e.g., *SQSTM1, BAX*, and *LAMP2*), consistent with the previous studies of *in vitro* TPC senescence (**Fig S6A**) (Schmid et al., 2019), raising the possibility of this pathway contributing to TPC survival.

**Figure 6.**
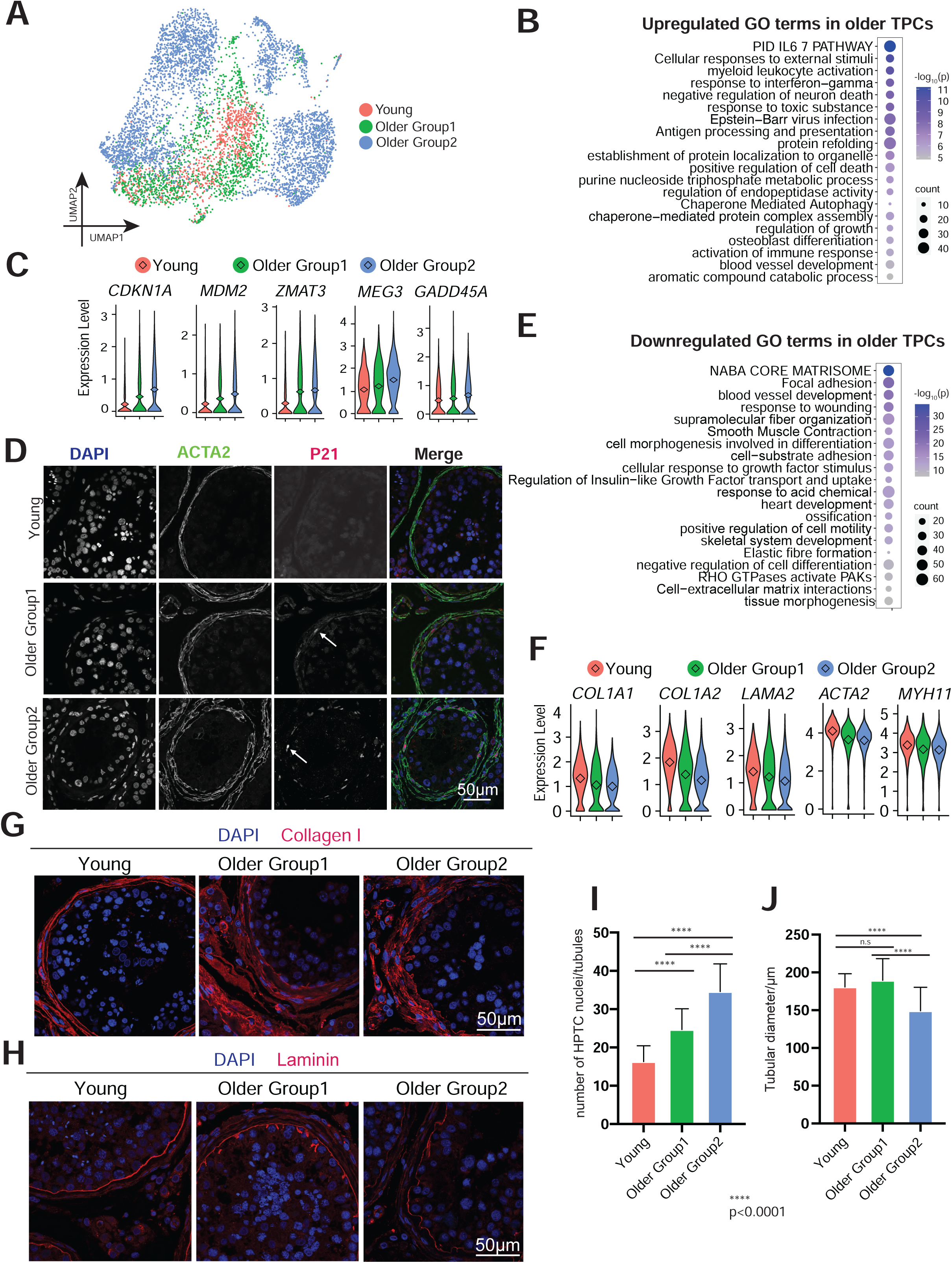
TPCs display morphological and functional alternations in older testes. (A) UMAP plot showing focused analysis of TPCs from Fig 1C. (B) Dot plot showing the top 20 upregulated GO terms enriched in the DEGs of older TPCs with P-value and gene numbers. (C) Violin plots with mean expression level highlighted inside showing upregulated DEGs of older TPCs. (D) Immunofluorescence images of P21 (encoded by *CDKN1A*, red) with an TPC marker, ACTA2 (green) in different groups, revealing increased P21 expression in older PTC cells. Nuclei were counterstained with DAPI (blue). Arrows indicate P21 positive cells. n = 6 human samples. (E) Dot plot showing the top 20 downregulated GO terms in older TPCs with P-value and gene numbers. (F) Violin plots showing downregulated DEGs of older TPCs. The diamond inside the violin plot represents mean. (G) Immunofluorescence images of Collagen I (red), revealing increased Collagen I deposition in older PTC cells. n = 6 human samples. (H) Immunofluorescence images of Laminin (red), revealing abnormal laminin deposition in older PTC cells. n = 6 human samples. (I) Quantification of the number of TPCs in cross-section of each seminiferous tubule in different groups, revealing a progressive increase in the numbers of TPCs around the walls of seminiferous tubules from Young through Older Group1 to Older Group2. Bars represent mean with SD of 20 independent tubules per group. n = 6 human samples. ****p < 0.0001, **p < 0.01 (two-tailed t-test). (J) Quantification of diameter of seminiferous tubules in different groups. Bars represent mean with SD of 20 independent tubules per group. n = 6 human samples. ****p < 0.0001, **p < 0.01 (two-tailed t-test).

Remarkably, core matrisome genes, namely *COL1A1/2, COL3A1, LAMA2/4*, and *LAMA4* were downregulated in older TPCs (**Fig 6E-F** and **S6C**). Considering that TPCs form the wall of seminiferous tubules, these results indicated defective integrity of the wall of seminiferous tubules. To test this, we stained collagen I and laminin (**Fig 6G-H** and **S6E-F**). In contrast to the mRNA data, collagen I protein surrounded the seminiferous tubules increased during aging and laminin showed an abnormal morphology with longer protrusions and irregular/unsmooth curve at the basement membrane in older testes. We further compared the downregulated genes of TPCs with published secreted proteomics data of *in vitro* senescence model of TPCs via mass spectrometry reported by Schmid et al (Schmid et al., 2019). Interestingly, of the 48 proteins detected downregulated in senescent TPCs, 11 proteins were likewise downregulated in our scRNA-seq data, with 8 of these 11 proteins functioning as key components of the wall of seminiferous tubules (**Fig S6D**). Taken together, the results indicate a decline of ECM secretion from older TPCs and an accumulation of ECM deposition, suggesting a complicated regulation of ECM in older testes.

Last, we observed GO terms related to the downregulation of smooth muscle contraction enriched in older TPCs (e.g., *ACTA2, MYH11, MYLK, MYL9*, and *TPM2*), indicating the ability to assist luminal transportation of sperm was impaired in older TPCs (**Fig 6E-F** and **S6C**). To validate, we induced replicative senescence in TPCs derived from independent donors, and tested TPCs for their ability to contract upon exposure to a known stimulus (FCS) (Schell et al., 2010). Both the time-lapse images and gel contraction assays demonstrated significant impairment in contraction with senescent TPCs (**Fig S6H-J**). Here, senescence of TPCs was confirmed by increased cell size and beta-galactosidase staining in advanced passage (**Fig S6G**), with cell viability in the gel contraction assay confirmed by the fluorescent dye Calcein AM (**Fig S6K**). Altogether, older TPCs show impaired integrity of the wall of seminiferous tubules and dysregulation of tubular transport of immobile spermatids.

### Spermatogenic patterns in older men are concordant with their BMI status

As Older Group1 and Older Group2 were markedly different in their physiology and genomics signatures, we probed available donor medical records for correlated factors that may affect their fertility. Surprisingly, all donors from Older Group1 (n=5) had low BMI (<27), while all donors (n=3) from Older Group2 had higher BMI (>30) (**Fig S7A**). Notably, donors within the Young group (which included both low and high BMI) all displayed normal and complete spermatogenesis irrespective of their BMI levels, which was further verified by checking the histology of 11 other young donors via PAS staining (**Fig S7B-C**). Taken together, our results suggest that aging itself confers a set of modest molecular changes that sensitize the testis for additional dysregulation – and when combined with the changes associated with obesity, may together lead to pronounced dysregulation, spermatogenic regression, and subfertility.

## DISCUSSION

Fertility rates decline in men as they age – however, the cellular and molecular basis of this decline is not understood, nor is it known whether lifestyle or environmental factors impact this decline(Wiener-Megnazi et al., 2012). To understand the cellular and molecular processes associated with aging, we performed scRNA-seq profiling of both young (17-22 years old) and aging/older adult (>60 years old) samples. The older donors were screened for having offspring as young adults, to ensure early-adult fertility. The profiles of our young samples were highly similar to each other. However, the older samples were clearly classified into two distinct groups: Group 1 displayed intact spermatogenesis and only weak/limited molecular signatures that distinguished them from young samples, whereas Group 2 displayed very limited spermatogenesis, which was accompanied by strong molecular signatures (discussed below) present within particular somatic cells of the testis – whereas the SSC populations of aging showed limited impact. Our findings on variation in spermatogenesis during aging are consistent with previous case-control cohort studies on spermatogenic capacity and regression in the older man (Paniagua et al., 1991; Pohl et al., 2021). Moreover, our work uncovered the first aging-associated molecular signatures – provided through deep scRNA-seq on all cell types of the testis – revealing misregulation of several pathways in key somatic cells of the testis niche, along with an apparent linkage of the phenotypic and molecular dysfunctions to age-associated elevated BMI.

### Aging men with elevated BMI are molecularly and phenotypically distinguishable

To determine whether differences between samples in Older Group1 and Group2 might relate to patient medical history, we examined all available donor information including BMI, smoking status and other components. Notably, elevated BMI emerged as a common factor: all donors from Older Group1 had BMI levels lower than 27, while all donors from Older Group2 had levels over 30. Importantly, our young donors did not display molecular signatures of aging or disrupted spermatogenesis irrespective of their BMI status. Previous studies have documented possible mechanisms underlying obesity-associated male subfertility, which largely involve changes in available hormone levels, the induction of inflammation, and increases of testicular temperature(Craig et al., 2017; Palmer et al., 2012). However, how aging and obesity can together exert synergistic problems on male fertility is largely unknown, especially at the molecular level. Here, we show that the aging-associated molecular signatures are greatly enhanced in the older men with high BMI and are associated with spermatogenic impairment. Therefore, we provide initial evidence that obesity may accelerate the effects of aging on male fertility and contribute to spermatogenic failure. Future studies will focus on examining at what age the impact of obesity on male fertility starts to emerge, and whether it is reversible by lowering BMI. Here, as BMI can be impacted by diet, exercise, and other factors (such as diabetes) – careful studies with large patient size will be needed to determine the causes that most impact fertility, which may lead to improved medical guidance for older men.

### Spermatogonial stem cell transcriptional signatures are similar in young and older men

Somewhat surprisingly, we find only subtle aging-related transcriptional changes within spermatogonia and spermatocytes and note two possible (and non-exclusive) reasons. First, germ cells that strongly deviate from the normal transcriptional program may undergo elimination in order to maintain the genetic or epigenetic fidelity of sperm and subsequent offspring. Second, undifferentiated spermatogonia are largely quiescent, with limited replication and metabolic activity, and therefore may represent a preserved pool of cells that has evolved to largely maintain fidelity over decades. This apparent lack of major impact on the spermatogonia has clinical implications, as improvements to the function of the somatic niche may then allow proper spermatogenesis from a healthy pool of spermatogonia. However, we note that distinguishable from young germ cells, aging germ cells demonstrate higher levels of DNA fragmentation, and display certain epigenetic changes (e.g., hyper DNA methylation) (Cao et al., 2020; Pohl et al., 2021). Consistent with the modest transcriptional changes in spermatogonia, we find no significant change in overall SSC transplant efficiency in both older groups, which also holds true in mice(Oatley and Brinster, 2012; Ryu et al., 2006). Notably, SSCs from older men show higher variation of colonizing potential in the xenotransplant assay, which is currently unexplained, but may be due to increased clonal amplification of selfish SSCs in older men (Maher et al., 2014), promoting further studies. Considering the dramatic aging-associated changes in niche cells we observed, it is highly likely that testicular niche undergoes aging-associated alterations that fail to keep SSC pool and development.

### Major aging-associated changes in somatic niche cells

We observe pronounced changes in aging testicular somatic cells, which may underlie diverse mechanisms of spermatogenic failure. First, the ‘nurse’ cells within the seminiferous tubule – the Sertoli cells – are known to display cell number reduction during aging, and undergo a series of morphological changes, such as oddly-shaped and large lysosomes, accumulation of lipid droplets, and mitochondrial metaplasia(Santiago et al., 2019). Our results validate the reduction of Sertoli cell number in aging men, and further suggest mechanisms that underly the morphological and cellular changes. For example, we reveal major metabolic changes, including dysregulation in lipid metabolic pathway as well as the decline of precursor metabolites and energy production, which may reduce nutrient support of germ cells, and limit germ cell numbers and progression. Second, through both genomics results and functional validation experiments we show that testosterone production by aging Leydig cells decreases, which can impair Sertoli cell activity and spermatogenesis.

Notably, in aging Leydig cells we observe transcriptional signatures of TPCs. This result can be understood by recognizing that human Leydig cells and TPCs are known to share a common progenitor prior to puberty, while these progenitor cells are not observed in the adult testis. This suggests that aging Leydig cells revisit earlier developmental transcriptional programs and decisions (Guo et al., 2020). In support of this possibility, we observe clear reductions in HH receptors in aging Leydig cells and the weakened HH signaling interaction of Sertoli-Leydig cells; results in keeping with prior *in vitro* studies that HH signaling induces the common progenitor cells to differentiate into Leydig cells in rats (Zhao et al., 2021). Third, we observe TPC hyperplasia in aging testes, which is associated with less contractility and abnormal secretion of basement membrane components, factors which could impair the transport of sperm along the tubule, as well as disrupting interactions between SSCs and the basement membrane. Here, further studies should focus on whether the increased number of TPCs involves compensation for their functional decline, and whether TPC hyperplasia may involve TPC proliferation.

In addition to cell type-specific alterations of somatic cells, we find upregulation of inflammation-induced genes shared across different testicular somatic cells. As a hallmark of aging, inflammation has been found in many types of organs(Azenabor et al., 2015; Ferrucci and Fabbri, 2018). The aging mouse testis also displays increased pro-inflammatory cytokines and hyperactive macrophages, and anti-inflammatory agents can increase the production of testosterone (Matzkin et al., 2016). Future work on identifying pro-inflammatory cytokines in aging human testis could pave the way for establishing anti-inflammatory treatments to improve aging-related fertility decline.

We also note limitations of the current study. Our work primarily focuses on transcriptomic profiling with additional protein validation; it would be interesting to explore the posttranscriptional and epigenetic mechanisms involved during testis aging. Furthermore, given the variations that appear to exist among different individuals, especially in older men, future studies designing to recruit large patient cohort could further validate and expand our findings regarding both transcription and transplantation. This may also allow us to better understand whether the link to BMI involves solely the index score, or is better correlated to diet, exercise, diabetes, or other contributing factors. Last, our study mostly focuses on the molecular and cellular in human testis; as aging also impacts the entire endocrine system and leads to altered hormone production, further studies should pay attention to examine the role of endocrine changes.

In summary, we profiled single cell transcriptomes of both germ cells and testicular niche cells from young and older human males, which serves as a foundational dataset for the scientific community. Further comparison analysis reveals the molecular changes taking place during testis aging, and provides potential candidate biomarkers for the diagnosis of aging-associated testis pathology. Our work also sheds light on how aging and other concurrent factors (such as BMI) may synergistically impact human testis functionality, which can provide guidelines to practice better lifestyle to improve fertility.

## ACKNOWLEDGEMENTS

We are grateful to the donors and their families. We thank Brian Dalley and Opal Allen in the HCI High-Throughput Genomics Shared Resource for sequencing expertise, Chris Stubben and Tim Parnell in the HCI Bioinformatics Share Resource for bioinformatics assistance, Xiang Wang in the Cell Imaging Cores of University of Utah, and DonorConnect staff for family consents and postnatal sample handling. This work was supported by NIA R01AG069725. Huntsman Cancer Institute core facilities was supported by NCI P30CA042014. B.R.C. is an HHMI investigator. J.B.S. was supported by the Swedish Childhood Cancer Foundation (TJ2020-0023). Studies with cultured TPCs were supported by Deutsche Forschungsgemeinschaft (DFG) project # 245169951, 427588170 and DFG MA1080/27-1. Fig 1A, 2G and 5G were created with BioRender.com.

## AUTHOR CONTRIBUTIONS

J.G. and B.R.C. conceived and supervised the project, and designed the experiments. J.G. and X.N. collected and processed donor samples. X.N. conducted computational analysis and validation experiments. K.I.A. and B.R.E. conducted testosterone ELISA assay with support from J.B.S. S.K.M. and M.S. conducted human spermatogonia xenotransplant assay supervised by K.E.O. N.S. and A.MI. performed studies with cultured TPCs, supervised by A.M. Sample acquisition was led by J.M.H. with input from B.R.C. and J.G. D.C. provided human testis samples. The manuscript was written by X.N. B.R.C. and J.G. with input and agreement of all authors.

## SUPPLEMENTAL FIGURE LEGENDS

**Figure S1.**
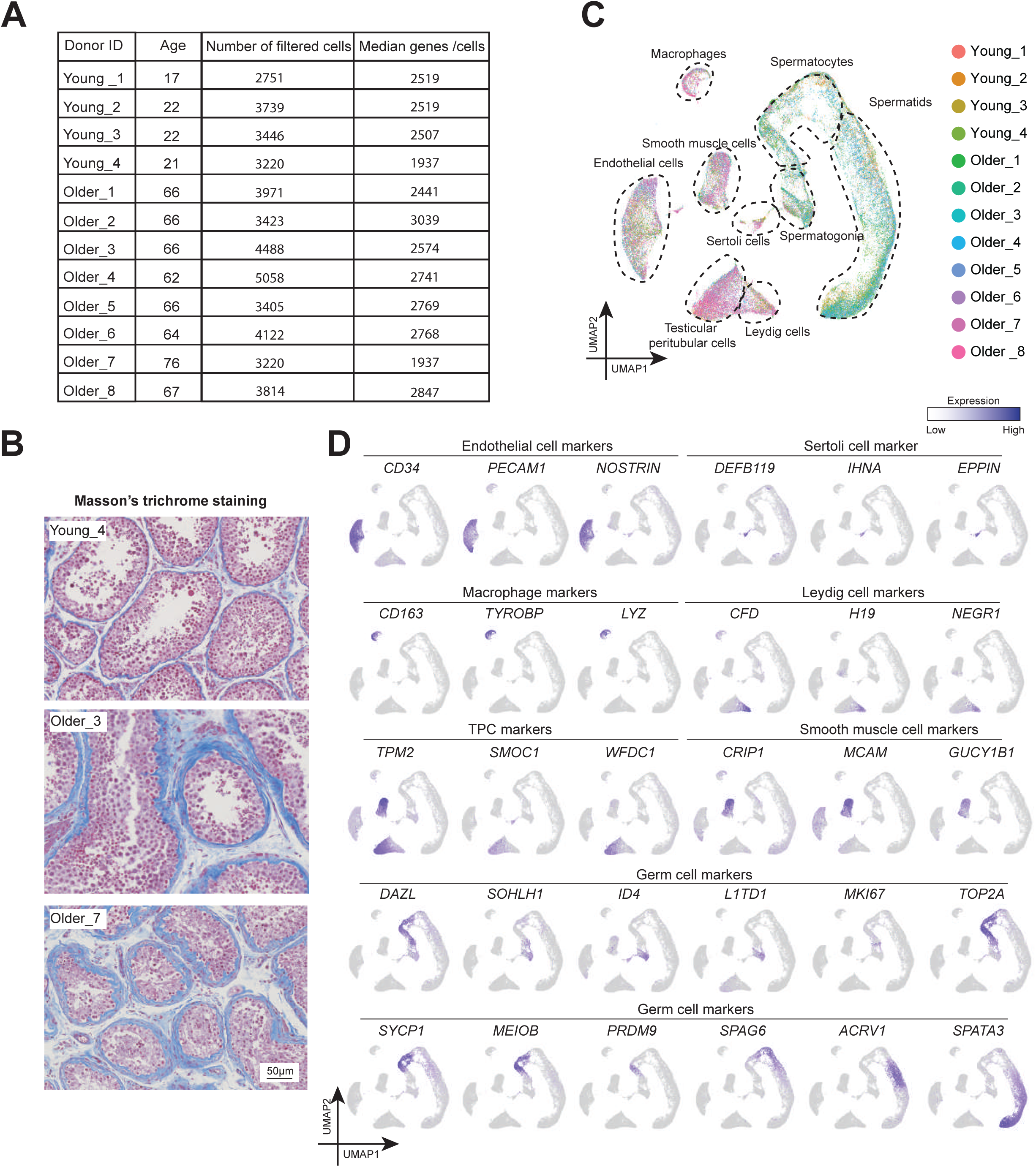
Information about donors and quality control of scRNA-seq. (A) Information summary of 12 donors sequenced in this study. (B) UMAP plot showing major testicular cell types with color according to the donors of origin. (C) Low magnification of the sections in Fig 1B, showing a thicker wall of seminiferous tubules and more ECM in interstitial tissues in older testis. The boxed regions represent the part of the area shown at the higher magnification in Fig 1B. (D) Expression of additional markers identifying major testicular cell types cast on the UMAP plot. Purple (or grey) represent high (or low) expression level as shown on the color key on top right.

**Figure S2.**
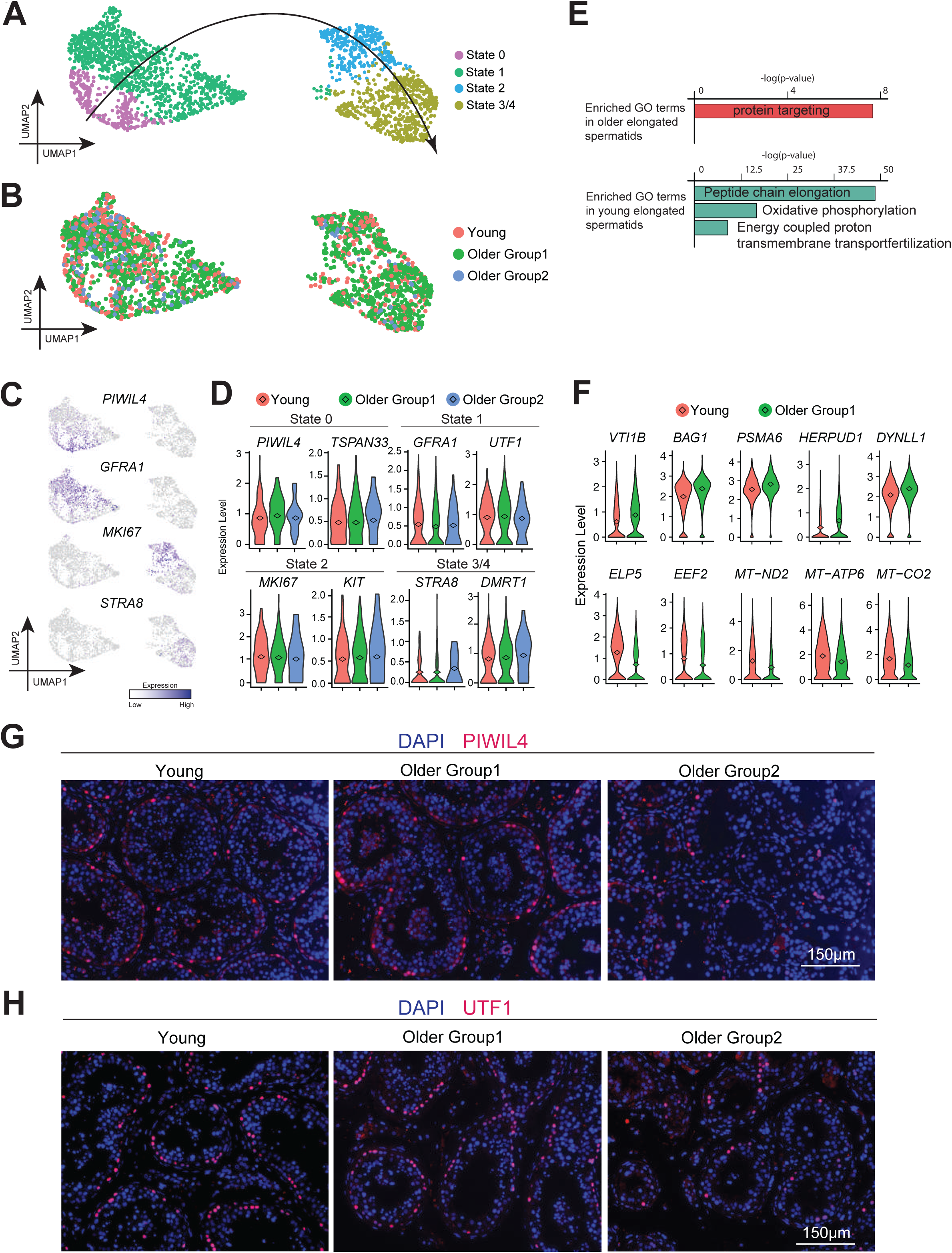
The transcriptome of human germ cells from older men shows high similarity with that of young men. (A) UMAP plot showing focused analysis of spermatogonia from Fig 2A. Cells are colored according to five discrete cellular states (States 0 to 4) as described in Guo et al., 2018. The arrow points to the development direction of spermatogonia. (B) UMAP plot showing focused analysis of spermatogonia from Fig 2A. Cells are colored according to Young versus Older Groups. (C) Expression of selected markers identifying major cellular states of spermatogonia cast on the UMAP plot. Purple (or grey) represents a high (or low) expression level. (D) Violin plot showing almost no difference among different groups for each state. The diamond inside the violin plot represents mean. (E) Bar plot shows upregulated (top) or downregulated (bottom) GO terms enriched in the DEGs of elongated spermatids when comparing Older Group1 to Young. (F) Violin plots showing upregulated DEGs of older elongated spermatids on the top panel and downregulated DEGs of older elongated spermatids at the bottom panel. The diamond inside the violin plot represents mean. (G) Low magnification of the sections in Fig 2E. Scale bar, 150 µm. (H) Low magnification of the sections in Fig 2F. Scale bar, 150 µm.

**Figure S3.**
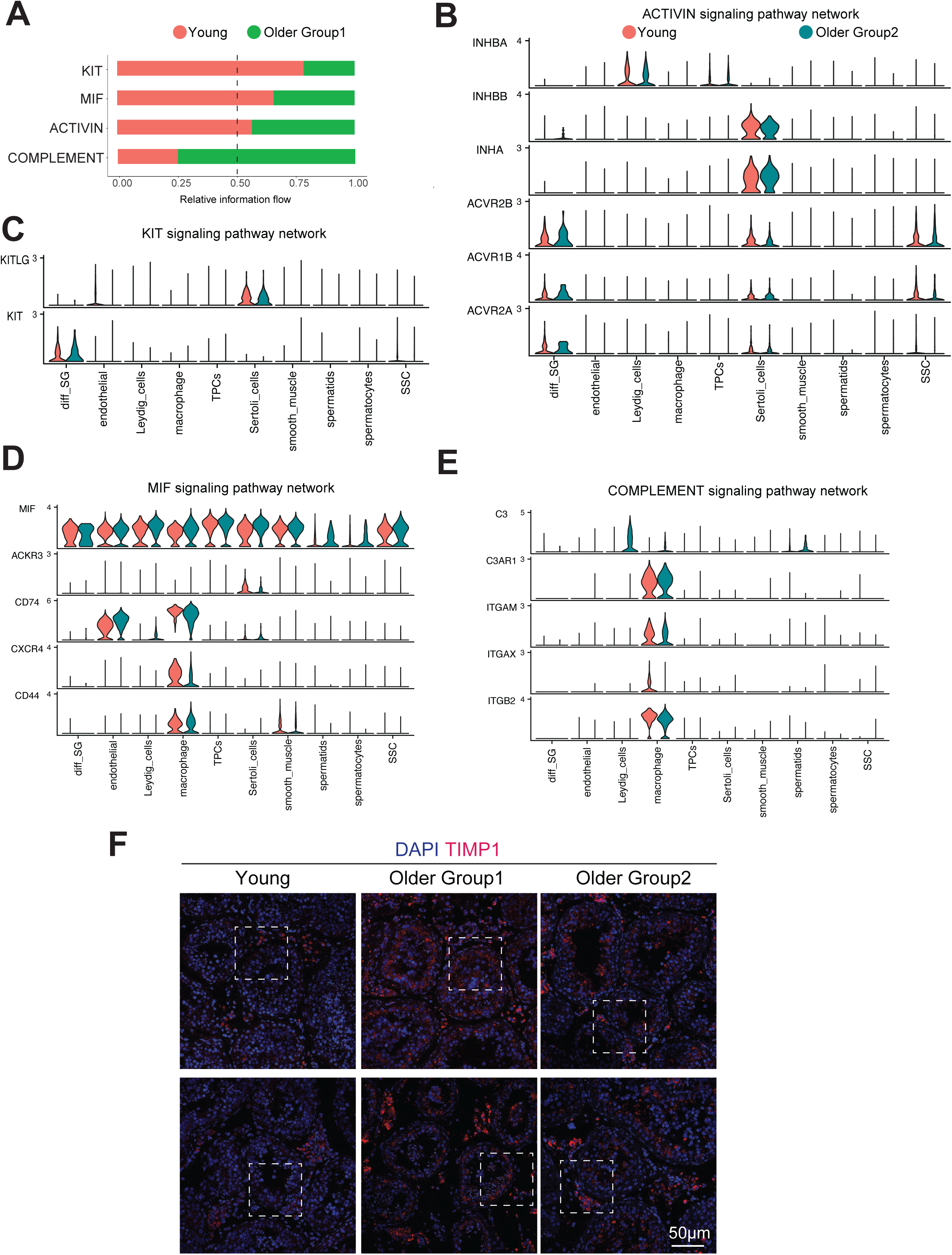
Differentially expressed signaling networks of cell-cell communication between young and older testis. (A) Bar graph (by CellChat analysis) illustrating representative information flow in Older Group1 (green) and Young (red). (B) Violin plot showing expression distribution of ACTIVIN signaling genes inferred by CellChat. (C) Violin plot showing expression distribution of KIT signaling genes inferred by CellChat. (D) Violin plot showing expression distribution of MIF signaling genes inferred by CellChat. (E) Violin plot showing expression distribution of COMLEMENT signaling genes inferred by CellChat. (F) Low magnification of the sections in Fig 3C. The boxed regions represent the part of the area shown at the higher magnification in Fig 3C.

**Figure S4.**
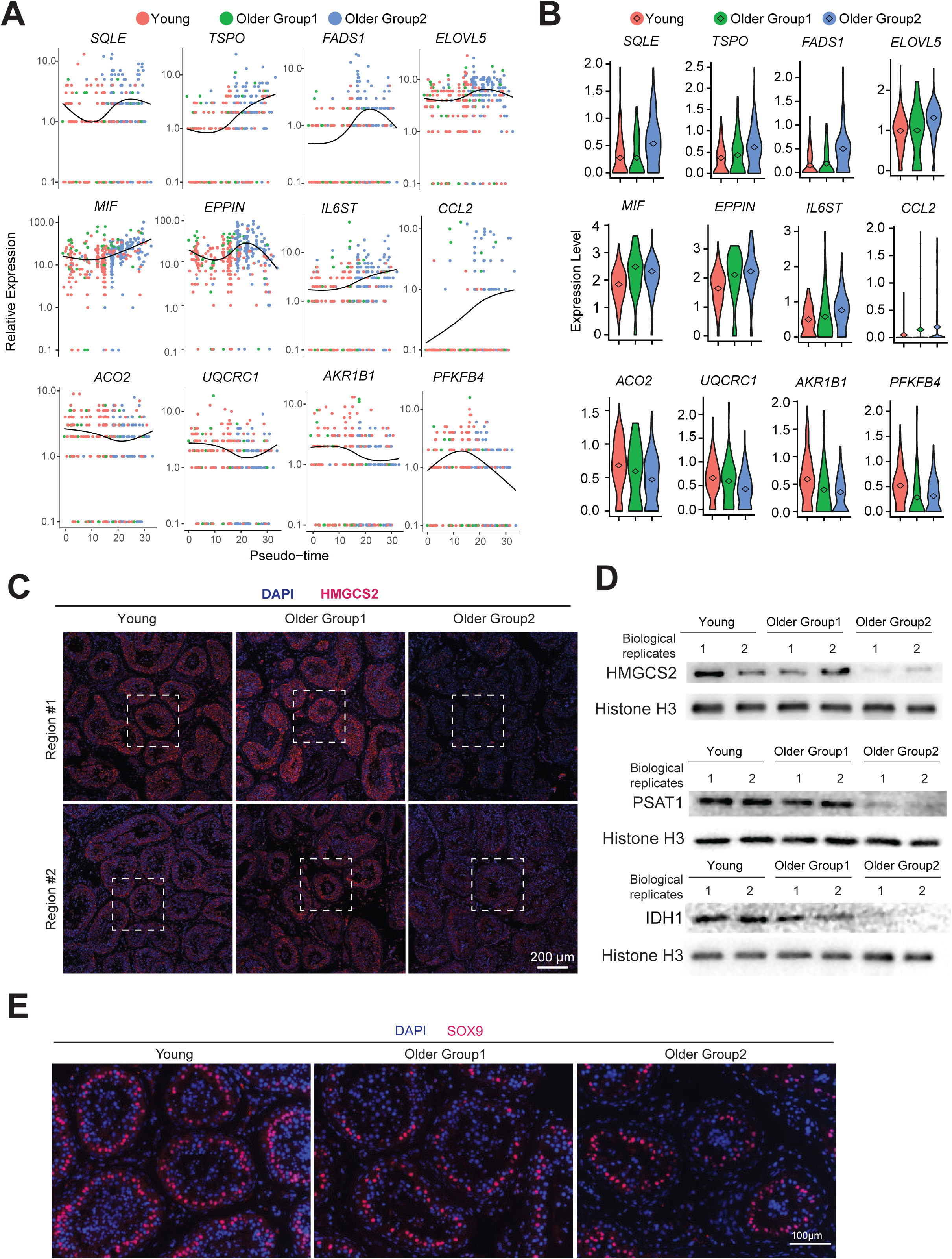
Gene expressions related to metabolism and inflammation in older Sertoli cells. (A) Expression levels of selected genes during Sertoli cell aging. The x-axis represents pseudotime as defined on Fig 4B, and the y-axis represents gene expression levels. (B) Violin plots showing expression levels of selected genes in Fig S4 during Sertoli cell aging. The diamond inside the violin plot represents mean. (C) Low magnification of the sections in Fig 4F. The boxed regions represent the part of the area shown at the higher magnification in Fig 4F. (D) Western blots of HMGCS2, PSAT1, and IDH1. Histone H3 was used for normalization. Total proteins were prepared from whole testicular tissues. Two biological replicates were used for each group. (E) Low magnification of the sections in Fig 4G.

**Figure S5.**
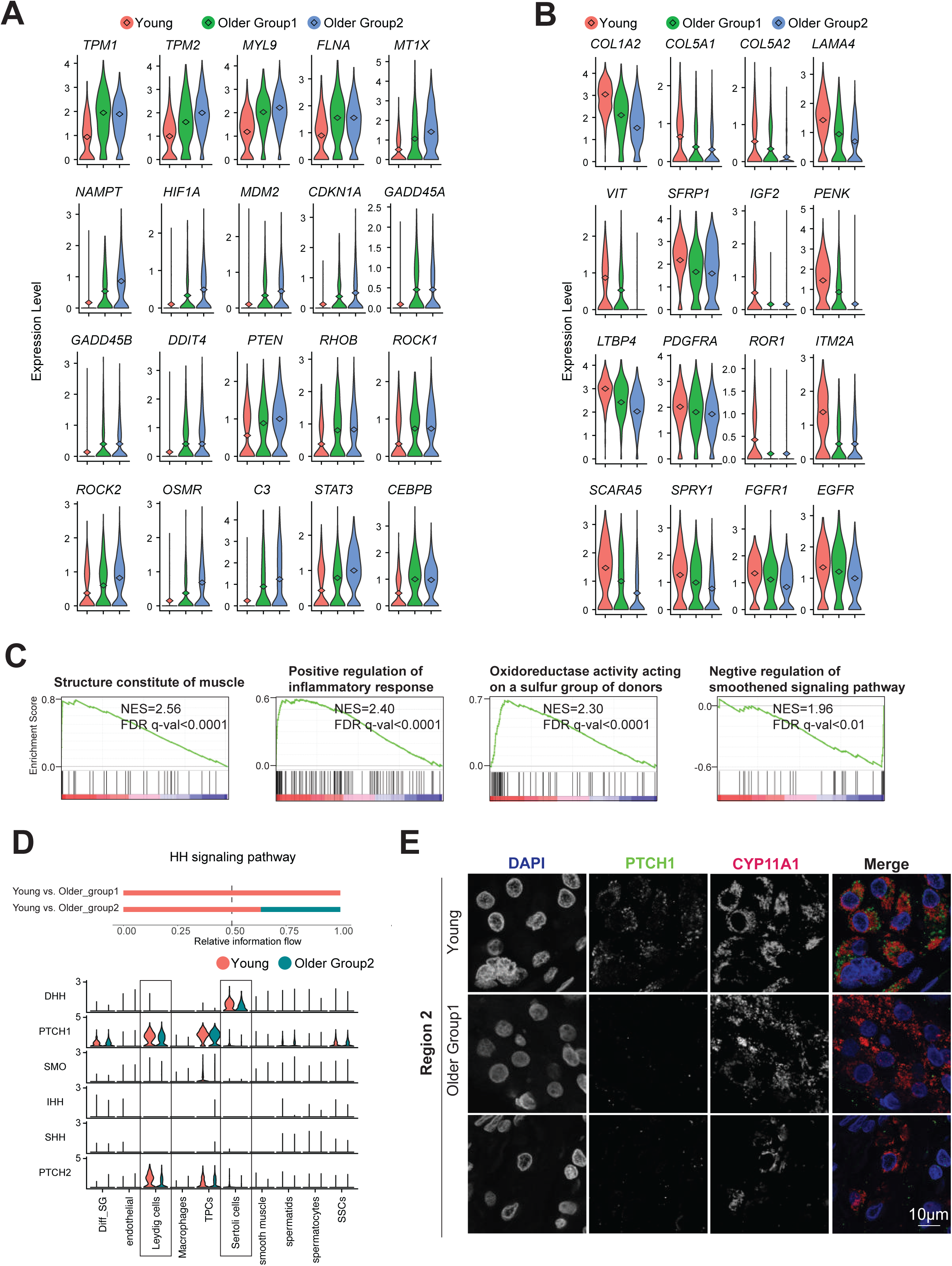
Signaling pathways are altered in older Leydig cells. (A) Violin plots showing upregulated DEGs of older Leydig cells. The diamond inside the violin plot represents mean. (B) Violin plots showing downregulated DEGs of older Leydig cells. The diamond inside the violin plot represents mean. (C) Gene Set Enrichment Analysis (GSEA) of Leydig cells in young and older testis. Representative enrichment plots are shown with significant GO terms on the top that correlated with older (red, left) or young (blue, right) Leydig cells. The peak of the green curves (enrichment score curve) on the red (left) side or the blue (right) side represent a positive or negative correlation with Older Leydig cells, respectively. Normalized enrichment score (NES) and Q-value are listed within each plot. (D) Top: Bar graph analyzed by CellChat illustrating representative information flow in Older and Young. Bottom: Violin plot showing expression distribution of HH signaling genes inferred by CellChat. (E) Independent experiments of Fig 5D using additional 3 human samples.

**Figure S6.**
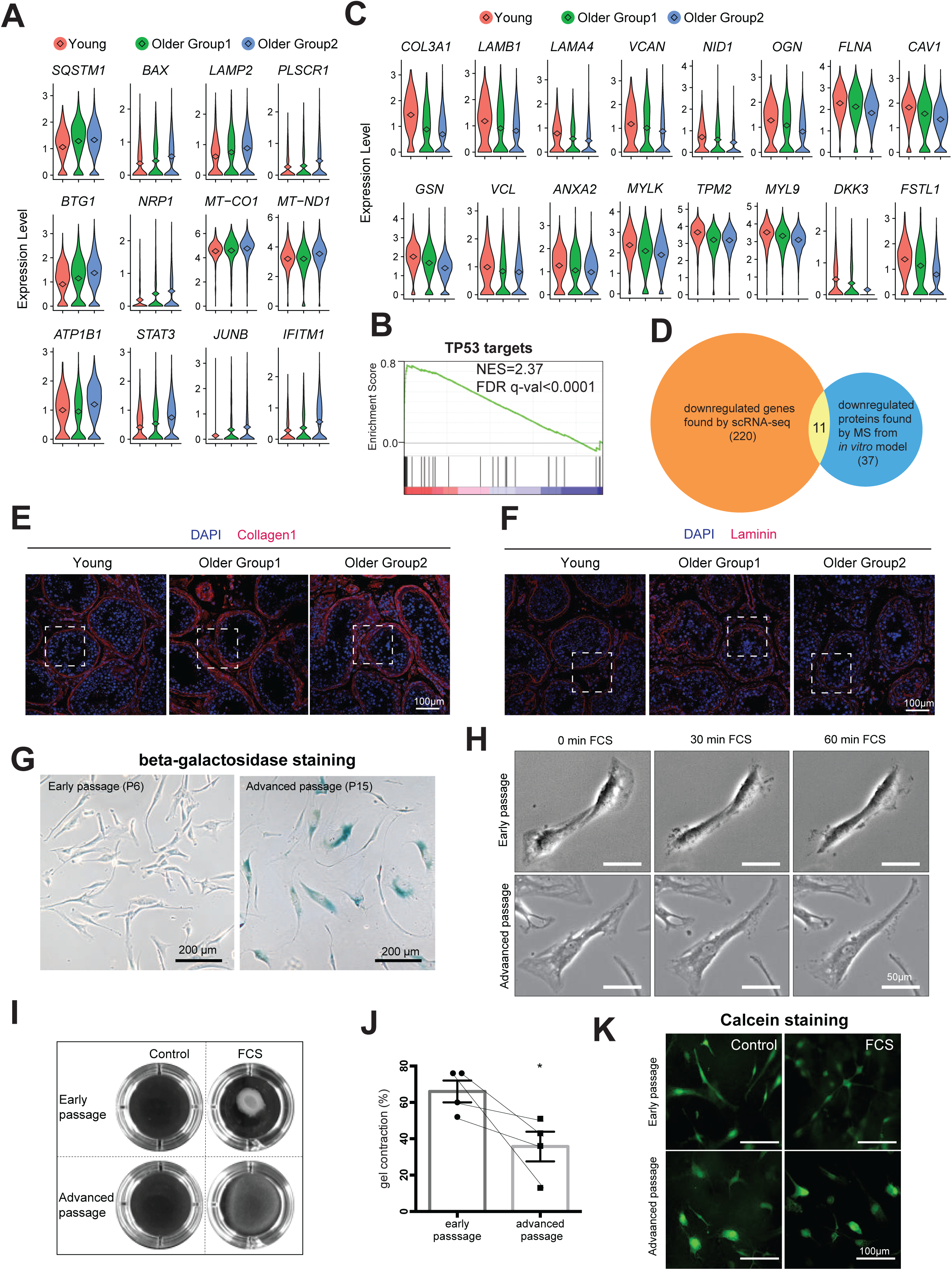
Disruption in ECM homeostasis in the testis of older men caused by TPCs and reduced contractile abilities. (A) Violin plots showing upregulated DEGs of older TPCs. The diamond inside the violin plot represents mean. (B) GSEA of TPCs in young and older testis. Representative enrichment plots are shown with significant GO terms on the top that correlated with older (red, left) or young (blue, right) TPCs. The peak of the green curves (enrichment score curve) on the red (left) side or the blue (right) side represent a positive or negative correlation with older TPCs, respectively. Normalized enrichment score (NES) and Q-value are listed within each plot. (C) Violin plots showing downregulated DEGs of older TPCs. The diamond inside the violin plot represents mean. (D) Venn diagram illustrates 11 shared genes/proteins downregulated in scRNA-seq of this study and secreted proteomics data of from in vitro senescence model of HPTCs via MS reported by Schmid et al. (E) Low magnification of the sections in Fig 6G. The boxed regions represent the part of the area shown at the higher magnification in Fig 4G. (F) Low magnification of the sections in Fig 6H. The boxed regions represent the part of the area shown at the higher magnification in Fig 4H. (G) Staining of senescence associated beta-galactodidase of cells in early (P6) and advanced passage (P15). Note that cell size was increased in most cells in P15. (H) Phase contrast micrographs of TPCs in early (above, P5) and advanced (below, P15) passages. Cells were monitored for 1 h during treatment with 30% FCS. A time dependent contraction of individual cells was observed. Considerable reductions of cell surface area are visible in cells in an early passage, in contrast to small changes of cells of an advanced passage. (I) Example of a collagen gel contraction assay for 24 h with TPCs of an early (P5) and advanced (P16) passage. Massive reduction of gel area of early passage after treatment with 30% FCS for 24 h, in advanced passage only slight reduction of the gel area is visible. Gel area remains nearly unchanged under control conditions. (J) Quantification of collagen gel contraction with TPCs after treatment with 30% FCS for 24 h (n = 4). Note massive reduction of gel area of early passages. In contrast, the corresponding cells in an advanced passage have a significantly reduced ability to contract. Bars represent mean with SD, statistical significance denoted with asterisk (* p < 0,05, t-test). Results from corresponding cells of different passages are connected with a line. (K) Example of evaluation of viability of cells in collagen gels directly after gel contraction assay using Calcein AM. Fluorescent cells were scored as alive.

**Figure S7.**
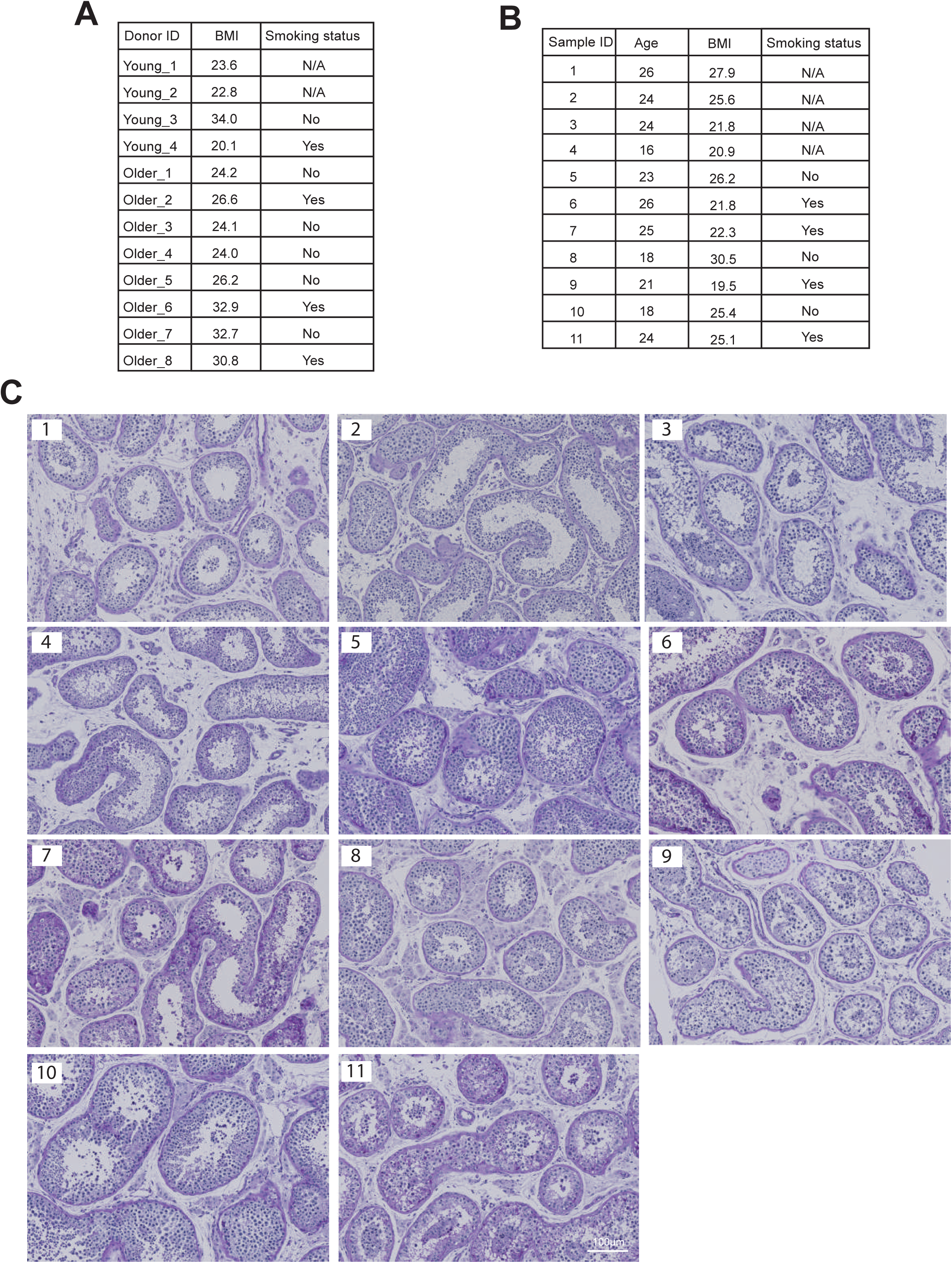
Testicular histology of donors with varying body mass index (BMI) and smoking status. (A) BMI and smoking status of donors who contributed to the single cell transcriptome profile in Fig 1A. (B) BMI and smoking status of 11 young donors other than donors contributing to the single cell transcriptome profile in Fig 1A. (C) Periodic acid–Schiff (PAS) staining of sections of testis from donors in Fig S7B.

## KEY RESOURCES TABLE

1 KEY RESOURCES TABLE
| REAGENT or RESOURCE | SOURCE | IDENTIFIER |
| --- | --- | --- |
| **Antibodies** |  |  |
| Rabbit anti-PIWIL4 | Invitrogen | Cat# PA5-31448, RRID AB_2548922 |
| Rabbit anti-UTF1 | Abcam | Cat# ab105090, RRID:AB_10903244 |
| Rabbit anti-DDX4 | Abcam | Cat# ab13840, RRID:AB_443012 |
| Mouse anti-TIMP1 [2A5] | Abcam | Cat# ab2464, RRID:AB_303083 |
| Rabbit anti-SOX9 | Abcam | Cat# ab185966, RRID:AB_2728660 |
| Rabbit anti-HMGCS2 | Sigma-Aldrich | Cat# HPA027423, RRID:AB_2672266 |
| Mouse anti-PTCH; clone OTI5C7 | Thermo Fisher Scientific | Cat# MA5-24893, RRID:AB_2725144 |
| Rabbit anti-CYP11A1 | Sigma-Aldrich | Cat# HPA016436, RRID:AB_1847423 |
| Mouse anti-p21 | Innovative Research | Cat# 33-7000, RRID:AB_87086 |
| Rabbit anti-ACTA2 | Abcam | Cat# ab5694, RRID:AB_2223021 |
| Rabbit anti-Collagen I | Abcam | Cat# ab34710, RRID:AB_731684 |
| Rabbit anti-Laminin | Abcam | Cat# ab11575, RRID:AB_298179 |
| Rabbit anti-PSAT1 | Proteintech | Cat# 10501-1-AP, RRID:AB_2172597 |
| Rabbit anti-IDH1 | Thermo Fisher Scientific | Cat# PA5-28206, RRID:AB_2545682 |
| Donkey anti-mouse Alexa Flour 488 | Invitrogen | Cat# A21202, RRID:AB_141607 |
| Donkey anti-rabbit Alexa Flour 594 | Invitrogen | Cat# A21207, RRID:AB_141637 |
| Goat anti-rabbit Alexa Flour 488 | Invitrogen | Cat# A32731, RRID:AB_2633280 |
| Rabbit anti-Histone H3 | Abcam | Cat# ab1791, RRID:AB_302613 |
| **Biological samples** |  |  |
| Human testis tissue | DonorConnect | N/A |
| **Chemicals, peptides, and recombinant proteins** |  |  |
| hCG | EmpowerR pharmacy | N/A |
| Busulfan | Sigma-Aldrich | Cat# 55-98-1 |
| **Critical commercial assays** |  |  |
| Chromium Next GEM Single Cell 3' GEM, Library & Gel Bead Kit v3.1 | 10 X genomics | Cat# PN-1000121 |
| Chromium Next GEM Chip G Single Cell Kit | 10 X genomics | Cat# PN-1000127 |
| **Deposited data** |  |  |
| Raw data files and processed data files for scRNA-seq | This paper |  |
| Experimental models: Cell lines |  |  |
| Human testicular peritubular cells | Technical University of Munich | N/A |
| Experimental models: Organisms/strains |  |  |
| Immune-deficient nude mice (NCr nu/nu) | Taconic | Cat# TAC:ncrnu, RRID:IMSR_TAC:ncrnu |
| Software and algorithms |  |  |
| Cell Ranger (version:4.0.0) | Zheng et al.,2017 | https://support.10xgenomics.com/single-cell-gene-expression/software/pipelines/latest/using/tutorial_ov |
| Seurat (version: 4.0.0) | Satija et al., 2015 | https://satijalab.org/seurat/index.html |
| GSEA (version: 3.0) | Tamayo, et al. 2005, Mootha, Lindgren, et al. 2003 | https://www.gsea-msigdb.org/gsea/index.jsp |
| metascape | Zhou et al. 2019 | https://metascape.org/gp/index.html#/main/step1 |
| monocle 2 | Qiu et al., 2017, Trapnell, et al. 2014 | http://cole-trapnell-lab.github.io/monocle-release/docs/ |
| ImageJ | NIH | https://imagej.nih.gov/ij/ |
| GraphPad Prism 8 | GraphPad Software Inc. | https://www.graphpad.com/scientific-software/prism/ |
| Leica LAS AF Lite | Leica | https://leica-las-af-lite.updatestar.com/ |

## MATERIALS AVAILABILITY

### Lead Contact

Further information and requests for reagents should be directed to and will be fulfilled by the Lead Contact, Jingtao Guo.

### Materials availability

This study did not generate new unique reagents.

### Data and code availability

All software tools can be found online (see Key resources table). The accession number for all sequencing data reported in this paper is GEO: GSE182786.

## EXPERIMENTAL MODEL AND SUBJECT DETAILS

### Human Testicular Tissues

All 12 human testicular tissues were obtained through DonorConnect. These sample were removed from deceased individuals who consented to organ donation for transplantation and research.

### Animals

All experiments utilizing animals in this study were approved by the Institutional Animal Care and Use Committees of Magee-Womens Research Institute and the University of Pittsburgh (assurance A3654-01) and was performed in accordance with the National Institutes of Health Guide for the care and use of laboratory animals.

### Human testicular peritubular cells (HTPCS)

Human testicular peritubular cells (HTPCS) were isolated from small testicular tissue fragments of patients with normal spermatogenesis and obstructive azoospermia, as described before(Albrecht et al., 2006; Schell et al., 2010; Walenta et al., 2018). All patients (age 28, 36, 38, 48 years) approved the use of their cells for scientific studies and the local ethical committee (Technical University of Munich, Department of Medicine, project number 169/18S) approved the study.

Cells were cultured at 37°C and 5% CO2 in DMEM High Glucose (Gibco) containing 10% FBS (fetal bovine serum; Capricorn Scientific) and 1% P/S (penicillin/streptomycin; Biochrom) and were propagated for several passages.

## METHOD DETAILS

### Sample Transportation and Storage

Upon surgical removal, pairs of whole testes were put in containers on ice for 1 to 2 hours of transportation. Then the tunica were removed and testicular tissues were cut into small pieces (∼500 mg to 1g each). 90% of tissues were directly transferred into cryovials (Corning) containing 1.5ml freezing medium (75% DMEM medium (Life Technologies) +10% DMSO (Sigma-Aldrich cat # D8779) + 15% fetal bovine serum (FBS) (Gibco)). The cryovials are placed in the isopropanol chamber (Thermo Fisher Scientific) and stored at –80°C overnight. Then the cryovials were transferred to liquid nitrogen for long-term storage.

### Sample Fixation

The remaining 10% testis tissues were fixed by 1X D-PBS (Thermo Fisher Scientific) containing 4% paraformaldehyde/PFA (Thermo Fisher Scientific) and agitated overnight at 4°C at 60 rpm. Fixed samples were then washed with cold PBS three times, and stored in PBS at 4°C until immunostaining or histochemistry processing.

### Sample Preparation for Single Cell RNA Sequencing

One cryovial of tissue was thawed quickly for each single-cell RNA sequencing experiment. Tissues were washed twice with PBS, and digested to single cell suspension as described previously. Briefly, tissues were scraped with razor blades and then digested with 1mg/ml collagenase type IV (Sigma Aldrich)/1mg/mL DNase I (Sigma Aldrich) and trypsin-ethylenediaminetetraacetic acid (EDTA; Invitrogen) /1mg/mL DNase I respectively at 37°C for 5 minutes. Then single cells are filtered through 40 um strainers (Thermo Fisher Scientific) and washed with D-PBS (Thermo Fisher Scientific). At last, cells were resuspended in D-PBS + 0.4% BSA (Thermo Fisher Scientific) at a concentration of ∼1,000 cells/uL ready for single-cell sequencing.

### Single Cell RNA-seq Performance, Library Preparation and Sequencing

The processes were instructed by the user guide of Chromium Next GEM Single Cell 3’ Reagent Kits v3.1 released by 10X genomics. Briefly, in the step of GEM Generation and Barcoding, cells were diluted for recovery of ∼5000 cells per lane and then loaded with master mix on Chromium Next GEM Chip G. After Post GEM-RT Cleanup, 12 cycles were used for cDNA Amplification. The resulting libraries were then sequenced on an Illumina Novaseq instruments with following setting: 28 cycles for Read 1, 10 cycles for i5 index, 10 cycles for i7 index, and 90 cycles for Read 2.

### Immunostaining of Testicular Tissues

Fixed tissues were embedded in paraffin and cut by 0.4 um per section. Sections were deparaffinized by Xylene and subsequently rehydrated in ethanol series (100%, 95%, 70%, 50%, water). Then, sections were incubated in sodium citrate buffers (pH 6.0) in a hot water bath (98°C) for 30 minutes for antigen retrieval. After blocking with blocking buffers (Thermo Scientific) for 30 minutes, slides were incubated with primary antibodies (1:200) which were diluted with Immunoreaction Enhancer Kit (Millipore sigma) overnight at 4°C in a humid chamber. The next day the slides were washed with PBS and incubated with secondary antibodies (1:400; Invitrogen) at room temperature for 2 hours in a humid chamber, followed by washes in PBS. Hoechst 33342 Solution was then diluted to 1:10000 to counterstain sections for 10 minutes. At last, the slides were washed with PBS again, mounted by Prolong Gold antifade mountant (Invitrogen), and sealed with nail polish ready to image. The antibodies are shown in the Key Resources Table.

### PAS Staining

Fixed tissues were embedded in paraffin and cut by 0.4 um per section. Sections were deparaffinized and hydrated to water. Then sections were oxidized in 0.5% Periodic Acid solution for 5 minutes, incubated in Schiff’s reagent for 15 minutes, and counterstained in Mayer’s hematoxylin for 1 minute. Between each step, rinse the sections in distilled water. At last, the sections were dehydrated and mounted using Xylene-based mounting media.

### Trichrome Staining

Fixed tissues were embedded in paraffin and cut by 0.4 um per section. Sections were deparaffinized and hydrated to water. Then sections were incubated sequentially in Weigert’s iron hematoxylin working solution for 10 minutes, in Biebrich scarlet-acid fuchsin solution for 10 minutes, and in phosphomolybdic-phosphotungstic acid solution for 15 minutes. Between each step, the slides were washed in distilled water. Then the sections were transferred directly (without rinse) to aniline blue solution and stained for 10 minutes. After washing briefly in distilled water, the sections were differentiated in 1% acetic acid solution for 5 minutes followed by rinsing in distilled water. At last, the sections were dehydrated and mounted using resinous mounting medium.

### Western Blot

Cryopreserved tissues were thawed quickly and homogenized by Dounce homogenizer with RIPA buffer. The lysates were further agitated for 30 minutes and centrifuged at 12,000g for 15 mins at 4°C. The supernatants were collected and protein concentration was calculated by Bradford Protein Assay (Bio-rad). The denatured lysates were electrophoresed on SDS-PAGE gels and transferred to polyvinylidene fluoride membranes (BioExpress). Then the membranes were blocked with 5% milk, incubated with primary antibodies overnight at 4°C, washed by TBST, and incubated with HRP-conjugated secondary antibodies for 1 hour at room temperature. Finally, signals were detected using ECL substrate (PerkinElmer). The antibodies are shown in the Key Resources Table.

### *Ex Vivo* Culture of Testicular Tissues and Testosterone Assay

The short-term tissue culture was adapted from a previous study. Briefly, the culture medium was KO-DMEM (Invitrogen) supplemented with 10% knock-out serum replacement (Invitrogen), and 1% penicillin/ streptomycin (Invitrogen). Cryopreserved tissues were thawed quickly and cut into small pieces (2∼4 mm^2^ in size), followed by weighing for statistical analysis. 3 pieces of tissues were immersed in 2ml medium per well of a 6-well plate. For each sample, 3 technique replicates were set, and 9 samples were tested in the experiments. The tissues were cultured at 34°C in 5% carbon dioxide for 24 hours and replaced with a fresh medium containing hCG and incubating for another 48 hours. Then, the supernatant ready for testosterone assay were collected by centrifuging the medium at 500 g for 5 minutes.

Testosterone concentrations were measured in culture media using the Roche e411 analyzer (Roche Diagnostics, USA) with the Elecsys Testosterone II enzyme-linked immunosorbent assay. Samples that exceeded the reportable range for the assay (>1500 ng/dL) were diluted and re-assayed according to manufacturer’s recommendations.

### Human to Mouse Xenotransplantation

Immune-deficient nude mice (NCr nu/nu; Taconic) were treated with chemotherapy agent busulfan (44mg/kg, Sigma) at 5-6weeks of age. Xenotransplantation was performed 5 weeks after busulfan treatment.

After tissue digestion, human testicular cells were suspended in MEMα medium (Thermo Fisher Scientific) containing 10% FBS (Thermo Fisher Scientific) and 10% trypan blue (Invitrogen). Approximately 7 μL of cell suspension was injected into the seminiferous tubules of a recipient mouse testis, using ultrasound-guided rete testis injections. Testes were recovered 8 weeks after transplantation, tunica removed, and seminiferous tubules dispersed with 1mg/mL Collagenase IV (Worthington) and 1mg/mL DNase I (Worthington) in D-PBS (Thermo Fisher Scientific). Tubules were fixed for 2hrs in 4% paraformaldehyde (Thermo Fisher Scientific) at 4°C and washed in D-PBS for 1hr (3-4X).

### Whole Mount Staining

After fixation, tubules were dehydrated in a series of graded methanol dilutions then incubated in a 4:1:1 ratio of MeOH: DMSO:H2O2v for 3 hours in 12-mm Trans well baskets (12-μm pore size; Corning Life Sciences). Tubules were then rehydrated in methanol dilutions and D-PBS, blocked with PBSMT blocking buffer (D-PBS, 0.02g/ml blotto dry milk powder & 10% Triton-X100) and stained with a rabbit anti-primate testis cell primary antibody (1:800) (Hermann et al., 2009) at 4°C overnight. Goat anti-rabbit Alexa Flour 488 (1:200; Invitrogen) was used to detect the primary antibody. Tubules were mounted on slides with Vectashield mounting medium containing DAPI with raised cover slips for fluorescence imaging.

### Senescence and Contractility of Human Testicular Peritubular Cells (HTPCs)

Cells from each patient were examined at an early passage (P5 – P7) and at an advanced passage (P15 – P17), i.e. when signs of senescence (increased cell size, cessation of proliferation) became visible, as described before(Schmid et al., 2019). To further document senescence, senescence-associated β-galactosidase staining was performed. To this end, HTPCs at P6 and P 15 were seeded onto glass cover slips and stained with a commercial kit (Senescence β-Galactosidase Staining Kit, Cell Signaling Technology) according to the manufacturer’s instructions and as described before(Schmid et al., 2019).Pictures were taken with a Zeiss Axiovert microscope (Zeiss GmbH).

Examination of contractility was employed by live cell monitoring and collagen gel contraction assays(Fleck et al., 2021; Schell et al., 2010).For live cell monitoring, cells were seeded onto imaging plates (35 mm µ-dishes; ibidi) and serum starved for 24 h. The cellular contractile response to 30 % FCS (in medium) was monitored over 60 min in a phase contrast microscope (Zeiss GmbH). In parallel control experiments medium without FCS was used.

For collagen gel contraction assays, 80.000 cells were embedded into collagen gel lattices (Collagen I-3D Gelling Kit, ScienCell) in a 24-well plate, coated with 0.2% BSA (Sigma-Aldrich). After 2 days of adjustment with daily medium changes, gel lattices were treated with or without 30 % FCS and pictures were taken after 24 h of stimulation. Collagen gel areas were evaluated with ImageJ and normalized to control conditions. In these experiments cell viability was assessed by staining with Calcein AM (2 µM; 15 min; Thermo Scientific) after treatment and pictures were taken with a fluorescence microscope (Leica).

### Microscopy

Immunofluorescence images were obtained through a Leica SP8 microscope. PAS and trichrome staining were scanned by a Zeiss Axioscan Slide Scanner. All raw images were processed by ImageJ. Slides of whole mount Immunofluorescence were observed under the Nikon Eclipse 90i microscope and analyzed using the NIS Elements Advanced Research software.

## QUANTIFICATIONS AND STATISTICAL ANALYSIS

### Processing of Single Cell RNA-seq Data

Cell Ranger v2.2.0 was used to demultiplex raw data by function *mkfastq*. The output Fastq files were then run with function *count* using default settings, including alignment (using STAR align to *GRCh38* human reference genomes), filtering, and UMI counting.

The generated UMI count matrices from 12 samples were loaded sequentially into R using the *Read10X* function of Seurat (http://satijalab.org/seurat/, R package, v.3). After adding the sample information in the row names of the matrices, all of them were merged and the Seurat object was created. According to the developer’s vignettes, the data were filtered and normalized. Specifically, cells were retained with >800 expressed genes and <50% of the reads mapped to the mitochondrial genome. The U-MAP and cluster analysis were performed on the combined dataset, using the top 5000 highly variable genes and 1-40 PCs. Cell type identification was also performed based on the tutorial. When analyzing each cell type separately, the cell barcodes of each cell type were extracted and the corresponding UMI count table from the total matrix was used for creating a new Seurat object.

The Seurat object of Sertoli cells was converted to CellDataSet object for importing into the *Monocle package* (v2.10.1). Pseudotime analysis was performed according to the default setting.

### Identification of Differentially Expressed Genes Associated with Aging

Aging_associated DEGs were found by the comparison of gene expressions from Young and old (both Old_group1 and Old_group2) using the FindMarkers function in Seurat. The Genes with expression level differences greater than 1.5 and P_adjust values less than 0.01 were set as the threshold. GO analysis of aging-associated DEGs was performed using the online tool *Metascope*.

To obtain the differentially expressed gene sets between young and old individuals, GSEA ranked all of the genes in the dataset based on differential expression. All parameters used to perform GSEA were default with C5 as database, 1000 permutations and “Signal2Noise” as ranking gene method.

### Analysis of Cell-Cell Communications

CellChat objects were created based on the UMI count matrix of each group(Young, Old_group1, and Old_group2) via *Cellchat* (https://github.com/sqjin/CellChat, R package, v.1). With “CellChatDB.human” set up as the ligand-receptor interaction database, cell-cell communication analysis was then performed via the default setting. The Comparation of a total number of interactions and interaction strength were obtained via merging the CellChat objects of each group by function *mergeCellChat*. The visualization of the differential number of interactions or interaction strength among different cell populations was achieved by function *netVisual_diffInteraction*. Last, differentially expressed signaling pathways were found by function *rankNet* and the signaling gene expression distribution between different datasets was visualized by function *plotGeneExpression*.

### Cell Counting, Tubular Diameter Measurement, SSC Colony Counting of Xenotransplantation and Statistical Analysis

For quantification of PIWIL4/UTF1/SOX9/ACTA2 expressing cells, single-positive cells were counted in cross-sections of round seminiferous tubules. 20 tubules were counted for each sample, and 2 samples were stained in each group.

For quantification of CYP11A1 expressing cells, 20 fields with round seminiferous tubule in the middle (0.04mm^2^/region) were counted for each sample, and 2 samples were stained in each group.

For quantification of the diameter of tubules, 20 roundish tubules were chosen from PAS staining for each sample, and 2 samples in each group were used. The shortest length of the tubules were measured as the diameter of the tubules by ImageJ.

For quantification of colonization activity in tubules, human spermatogonia colonies were counted if colonies were located on the basement membrane of the mouse seminiferous tubules, contained at least 4 cells in a continuous area (≤ 100µm between cells), were ovoid-shaped and had a high nuclear to cytoplasmic ratio.

The bars represent means ± SD of independent tubules. Data were analyzed using an unpaired two-sided Student’s t-test by PRISM 8 Software. p < 0.05 was considered statistically significant.

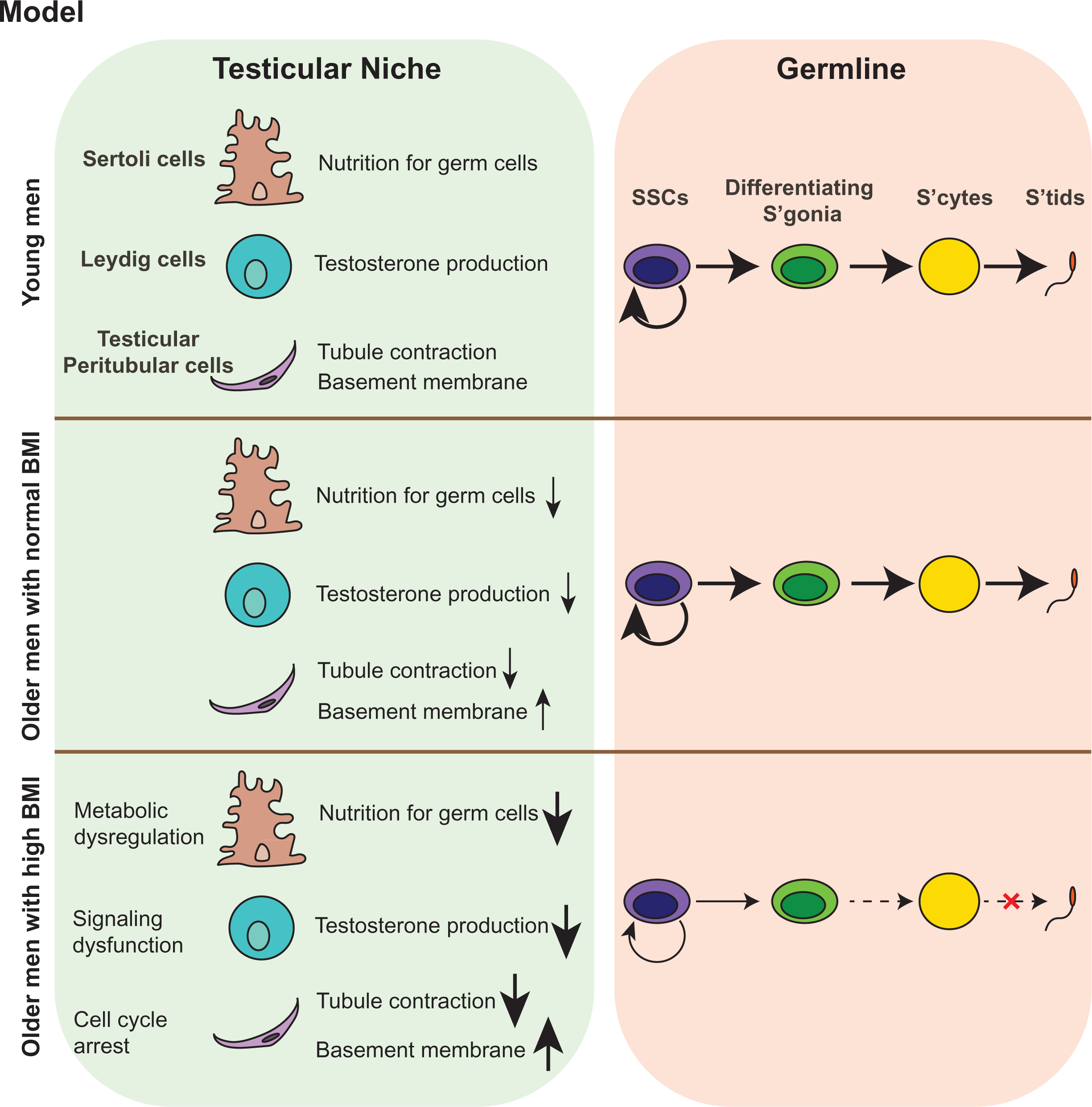

